# Endothelial Prolyl Hydroxylase 3 Mitigates Maladaptive Inflammation to Promote Post-Ischemic Kidney Repair

**DOI:** 10.1101/2025.04.21.649850

**Authors:** Rajni Sharma, Ratnakar Tiwari, James O’Sullivan, Gabriella S. Borkowski, Si Young An, Edward B. Thorp, Navdeep S. Chandel, Susan E. Quaggin, Pinelopi P. Kapitsinou

**Author notes:** Address correspondence and lead contact: Dr. Pinelopi P. Kapitsinou, Division of Nephrology and Feinberg Cardiovascular and Renal Research Institute, Feinberg School of Medicine, Northwestern University, 303 East Superior Street, SQBRC 8-408, Chicago, IL 60611. Equal Contribution (Rajni Sharma, Ratnakar Tiwari).

## Abstract

Acute kidney injury (AKI) is a major health concern and well-established risk factor for the development of chronic kidney disease (CKD). Tissue hypoxia is a prominent feature of the injured kidney that shapes the biological behavior of parenchymal and immune cells. Endothelial cells have important immunomodulatory roles, but the impact of dysregulated oxygen sensing on their responses remains poorly understood. By leveraging a combination of conditional mouse strains, single-cell analyses of mouse and human samples, and in vitro experiments, we demonstrate that the oxygen sensor PHD3 in the endothelium suppresses maladaptive inflammatory responses, thereby promoting kidney repair. Specifically, post-ischemic inactivation of endothelial PHD3, but not PHD1, leads to maladaptive kidney repair, characterized by increased fibrosis and inflammation. scRNA-seq analysis of the postischemic endothelial PHD3-haplodeficient kidney shows an endothelial IFN-γ gene signature, resembling the responses seen in patients with severe AKI. By performing loss- and gain-of-function experiments in vitro, we demonstrate that PHD3 regulates IFN-γ responsive pro-inflammatory signatures in a HIF-dependent manner. Consistent with this, simultaneous deletion of ARNT in endothelial PHD3-deficient mice restored kidney repair. Thus, our findings provide novel mechanistic insights into endothelial oxygen sensing and the AKI-to-CKD transition, highlighting potential therapeutic avenues for mitigating disease progression.

**TRANSLATIONAL STATEMENT:** Using a genetic approach, we identified endothelial PHD3 as a critical regulator of kidney repair. Post-ischemic loss of endothelial PHD3—but not PHD1—led to maladaptive outcomes marked by increased injury, fibrosis, and inflammation. Mechanistically, we show that PHD3 restrains IFN-γ–driven pro-inflammatory signaling through a HIF-dependent pathway. These findings reveal a central role for endothelial oxygen sensing in dictating kidney recovery and underscore the importance of cellular context in responses to hypoxia. Given the expanding clinical use of non-selective PHD inhibitors, our work highlights the need for more targeted, cell-specific approaches to modulate hypoxia pathways in kidney disease.

## INTRODUCTION

Every year, acute kidney injury (AKI) leads to 2 million deaths worldwide, affecting patients suffering from ischemia caused by surgery, sepsis, trauma, or toxic drug reactions ^1,2^. AKI often leads to chronic kidney disease (CKD) ^3,4^, which is projected to become the fifth leading cause of death by 2040 ^5^. Nevertheless, there are currently no effective treatments for AKI, highlighting the urgent need for novel preventive and therapeutic strategies.

The kidney has a remarkable capacity for repair following injury ^6^. However, when tissue injury is severe, repetitive, or the wound-healing process is disrupted, maladaptive responses lead to irreversible fibrosis ^7,8^. Inflammation is a central feature of maladaptive repair, involving complex interactions between kidney structural cells and the immune system ^9–11^. In particular, endothelial cells (EC) have emerged as key immune regulators, controlling the entry of innate and adaptive immune cells into tissues by expressing molecules such as adhesion proteins, and chemokines ^12^. Following AKI, EC are well equipped to sense cues and changes within the tissue microenvironment, such as hypoxia and cytokines, which influence their role in shaping immune responses ^13^. Persistent endothelial dysfunction leads to capillary rarefaction, impairing oxygen and nutrient delivery to the kidney tissue, and promoting CKD progression ^14,15^.

The prolyl-4-hydroxylase domain-containing proteins (PHD1–3, also known as EGLN1-3) act as molecular oxygen sensors, orchestrating cellular adaptation to hypoxia by regulating hypoxia-inducible factor (HIF)-α protein stability ^16^. In normoxia, PHDs hydroxylate the HIF-α subunit, targeting it for ubiquitin-proteasome-mediated degradation ^16^. Under hypoxic conditions, reduced hydroxylation allows HIF-α to escape ubiquitylation and undergo nuclear translocation, where it heterodimerizes with the constitutively expressed HIF-β or aryl hydrocarbon receptor nuclear translocator (ARNT) subunit. The active heterodimer can then recognize and bind to specific genomic DNA-binding sites (HREs; hypoxia response elements) and activate transcriptional programs that control erythropoiesis, angiogenesis, and metabolism ^16,17^. Among these isoforms, PHD2 (EGLN1) serves as the primary HIF prolyl hydroxylase, predominantly regulating HIF1α levels during normoxia ^18^. In contrast, PHD1 and PHD3 play more context-dependent roles, influenced by their distinct tissue-specific expression patterns ^19^.

In the kidney, HIFs drive cell type-specific responses essential to both homeostasis and disease, yet the roles of their upstream regulators, PHDs, remain incompletely understood. We recently reported that while endothelial PHD2 deficiency alone induced after ischemic AKI does not influence kidney repair outcomes, the combined loss of PHD1, PHD2, and PHD3 significantly worsens renal fibrosis and inflammation ^20^. These findings suggest that endothelial oxygen sensing regulates kidney repair through mechanisms involving the broader PHD family, highlighting the need to define the specific roles of endothelial PHD isoforms. Here, we employed a genetic approach using multiple endothelial-specific, inducible knockout mouse models and complementary in vitro studies to investigate the role of oxygen sensors PHD1 and PHD3 alone or in conjunction with ARNT in post-ischemic kidney repair.

## METHODS

### Animals

The generation and genotyping of mice with floxed alleles for *Phd1* (*Egln2*), *Phd3* (*Egln3*), and *Arnt* have been described previously ^21,22^. For PHD immunostaining studies, we generated *Cdh5-Cre; mT/mG* mice by crossing *Cdh5-Cre* with *ROSA26-ACTB-tdTomato,-enhanced green fluorescent protein* (*-EGFP*) reporter mice (JAX stock #007576), as previously reported ^23^. For inducible endothelial-specific inactivation of the floxed alleles, we used the *Cdh5(PAC)-CreERT2* transgenic mouse line ^24^. Cre-mediated inactivation was induced by four intraperitoneal (i.p.) injections of tamoxifen dissolved in 10% ethanol in corn oil at 20 mg/mL (3 mg/mouse), given every other day. Male mice aged 8–10 weeks underwent unilateral renal ischemia-reperfusion injury (uIRI) surgery as previously described ^20^. All animal studies were conducted in accordance with the NIH guidelines for the use and care of live animals and were approved by the Institutional Animal Care and Use Committee at Northwestern University.

### Preparation of single-cell suspension

Single-cell suspensions were generated using the Multi Tissue Dissociation Kit 2 (Miltenyi Biotec, Cat# 130-110-203), following the manufacturer’s protocol.

### scRNA-seq library generation and sequencing

Cells were loaded into the Chromium Controller (10X Genomics, Cat# PN-120223) using a Chromium Next GEM Chip G (10X Genomics, Cat# PN-1000120) to generate single-cell gel beads in emulsion (GEM), following the manufacturer’s protocol. cDNA synthesis and library preparation were carried out using the Chromium Next GEM Single Cell 3’ Reagent Kits v3.1 (10X Genomics, Cat# PN-1000286) and the Single Index Kit T Set A (10X Genomics, Cat# PN-1000213) according to the provided guidelines. The cell hashing library was prepared according to BioLegend’s protocol for HTO library construction using single index adapters. Library quality was assessed using the Agilent Bioanalyzer High Sensitivity DNA Kit (Agilent Technologies, Cat# 5067-4626), and quantitative analysis was performed with the Qubit DNA HS Assay Kit. The pooled multiplexed libraries were sequenced on an Illumina HiSeq 4000 using 2×50 paired-end kits, with Read1 (28 bp) capturing cell barcodes and UMI and Read2 (90 bp) sequencing transcripts. Raw sequencing data in base call format (. bcl) was demultiplexed and converted to FASTQ format using Cell Ranger (10X Genomics), which was also used for read alignment to the mouse reference genome (mm10) and for gene expression quantification.

### Statistical analysis

Data are reported as mean values ± SEM. GraphPad Prism 9 (GraphPad Software, La Jolla, CA, USA) was used for the statistical analyses. Specifically, two-group comparisons were performed using unpaired Student’s t-test with Welch’s correction, whereas multi-group comparisons were performed using one-way analysis of variance (ANOVA) with Sidak’s multiple comparison test. A two-sided significance level of 5% was considered statistically significant.

## RESULTS

### PHD isoforms demonstrate distinct, time-dependent expression in the kidney endothelium following AKI

To gain insights into the role of PHDs within the renal vasculature, we analyzed publicly accessible snRNA-seq data from mouse kidney IRI studies (Figure 1a) ^25^. Focusing on the EC cluster, we observed a distinct pattern of PHDs expression in response to AKI. Specifically, *Phd1* expression increased early following IRI (4 h and 2 days post-IRI) and returned to baseline at later time points. In contrast, PHD3 was downregulated 4 h post-IRI, followed by an increase starting at 12 h, which subsequently persisted. PHD2 showed low expression levels under baseline conditions without robust changes following IRI. Next, we examined the expression pattern of PHDs using publicly available single-nucleus RNA sequencing (snRNA-seq) data from kidney tissues of patients with severe AKI ^26^. Consistent with the mouse studies, human kidney EC from AKI patients showed increased expression of *PHD3* compared to controls (Supplemental Figure 1). On the other hand, *PHD1* mRNA was not detected, whereas *PHD2* expression increased in AKI (Supplemental Figure 1).

**Figure 1.**
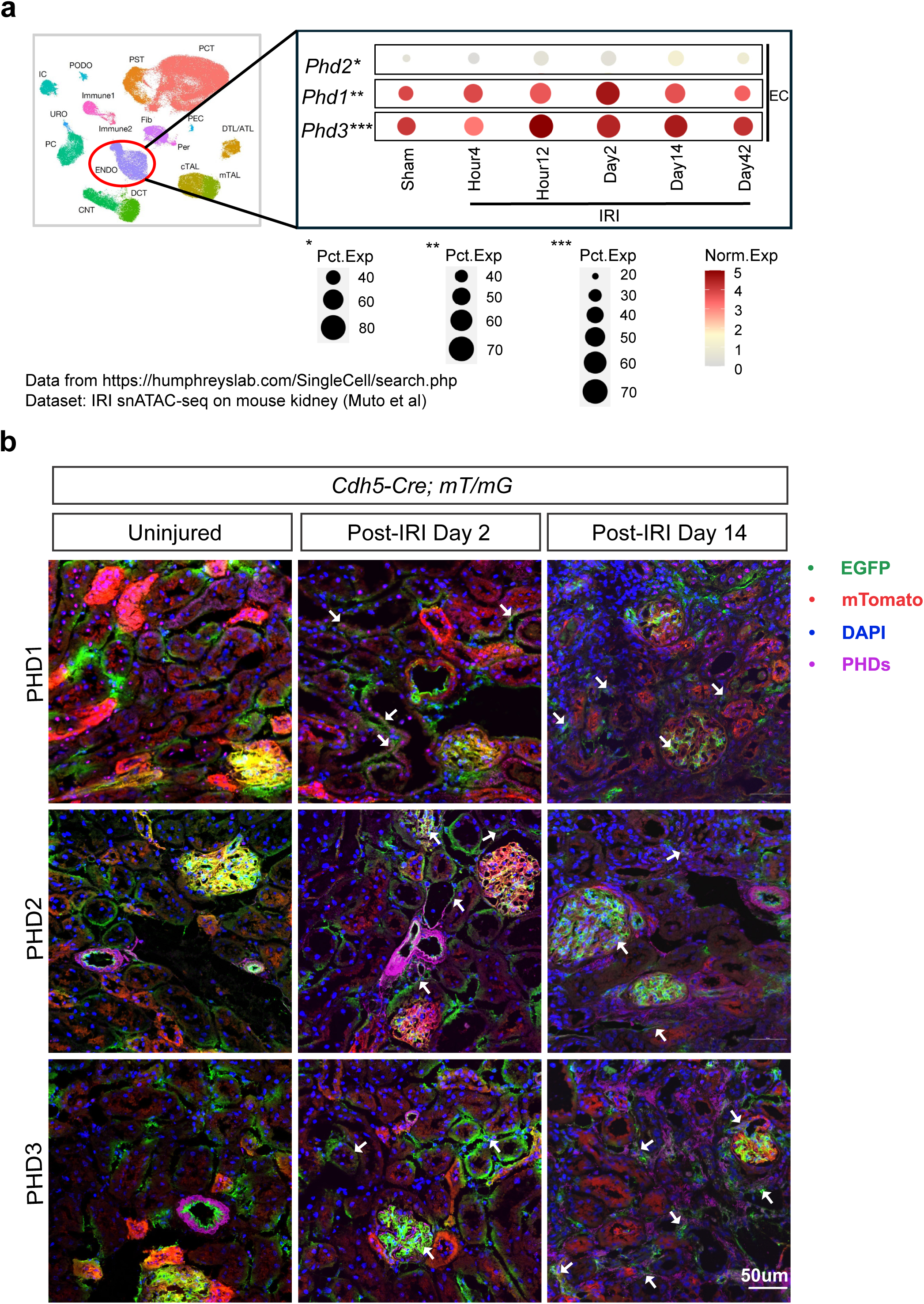
Differential expression of PHD1, PHD2, and PHD3 in murine kidney endothelium following ischemic kidney injury. **(a)** Dot plots showing expression of *Phd1* (*Egln2*), *Phd2* (*Egln1*), and *Phd3* (*Egln3*) transcripts in EC clusters from kidneys of mice subjected to bilateral IRI and analyzed at 4 h, 12 h, 2 days, 14 days, and 6 weeks post IRI. The expression levels of *PHD* genes were extracted from the online interface of Humphreys Lab (https://humphreyslab.com/SingleCell/displaycharts.php) using data derived from Muto et al. (25). **(b)** Representative images of immunofluorescence staining for PHD1, PHD2, and PHD3 (magenta) in uninjured and post uIRI kidney sections from *Cdh5-Cre; mT/mG* reporter mice at the indicated experimental time points. Non-recombined cells express membrane-bound mTomato (red), while recombined cells express membrane-bound EGFP (green). Individual channels are shown in Supplemental Figure 2. IRI, ischemia-reperfusion injury. Scale bar indicates 50 μm. IRI, ischemia reperfusion injury

To examine whether transcriptional changes reflected endothelial PHD protein expression, we performed immunohistochemistry on kidneys from *Cdh5-Cre; mT/mG* mice after unilateral IRI (uIRI), as previously described ^27^. Our study included two time points following uIRI, days 2 and 14, whereas uninjured kidneys were used as a baseline for comparison. In contrast to PHD1 and PHD2, which were not markedly changed at the selected time points, PHD3 showed increased expression, particularly on day 14 post-uIRI (Figure 1b and Supplemental Figure 2). Taken together, our findings highlight the distinct, time-dependent expression of PHD isoforms in response to AKI, with a delayed but sustained increase in PHD3 following AKI in both mice and humans.

### Post-ischemic inactivation of endothelial PHD3 but not PHD1 exacerbates post ischemic kidney injury and promotes maladaptive kidney repair

To better understand the roles of endothelial PHD1 and PHD3 in AKI, we performed genetic studies in which these isoforms were individually inactivated in the endothelium following kidney uIRI. First, we generated *Cdh5-CreER^T2^; Phd1^f/^*^f^ (*PHD1^iEC^*) mice by crossing mice homozygous for a conditional *Phd1* allele with *Cdh5-CreER^T2^* transgenic mice. Adult *PHD1^iEC^* mice and their *Cre-*littermates were subjected to 25 min of uIRI, followed by four i.p. tamoxifen injections and analysis on day 14, as shown in Figure 2a. Tubular injury assessment of H&E-stained sections from day 14 post-IRI kidneys revealed that *PHD1^iEC^*had comparable tubular damage to control mice. Quantitative analysis of Picro-Sirius red staining showed no significant difference in collagen deposition between the *PHD1^iEC^* mutants and controls (n=7-8) (Figure 2b). Although day 14 post-ischemic kidneys showed significantly increased transcript levels of pro-fibrotic genes, lysyl oxidase like2 (*Loxl2*), transforming growth factor beta 1 (*Tgfb1*), smooth muscle cell a-actin (*Acta2*), and hepatitis A virus cellular receptor 1 (*Havcr1*, gene encoding KIM-1) compared to their corresponding contralateral kidneys, no significant change was observed between the two genotypes (n=7) (Figure 2c).

**Figure 2.**
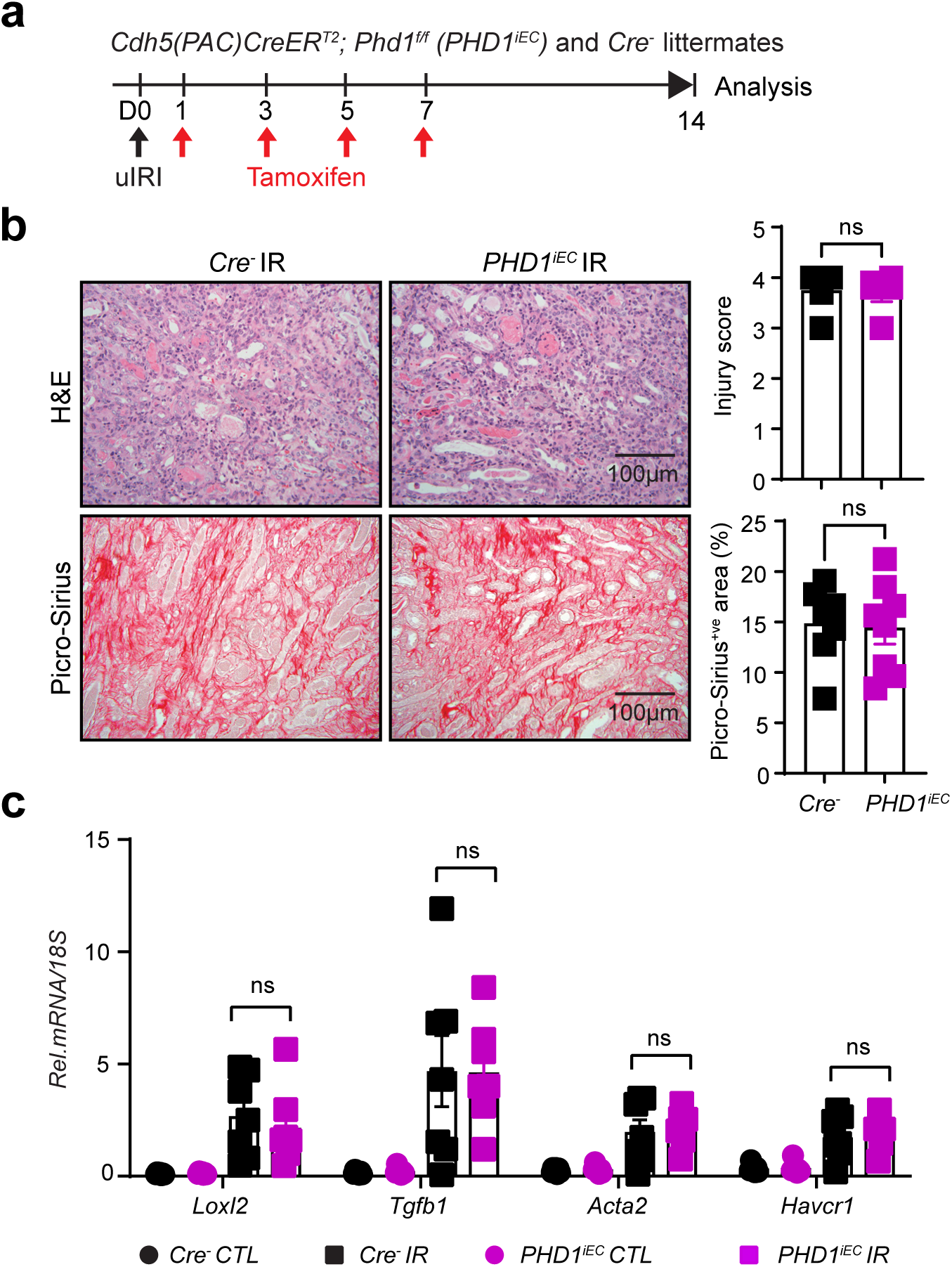
Inactivation of endothelial PHD1 after IRI does not affect post-ischemic kidney fibrosis. **(a)** Scheme illustrating the experimental strategy applied for renal uIRI studies. *PHD1^iEC^* mice and their *Cre-* littermates were subjected to 25 min of unilateral renal artery clamping. Treatment with tamoxifen started on day 1 post-uIRI, involving 4 i.p. doses administered every other day. Mice were sacrificed for histopathological and molecular analysis on day 14 post-uIRI. **(b)** Representative images of H&E- and Picro-Sirius red-stained sections from day 14 postischemic kidneys of *PHD1^iEC^* mutants and their *Cre-* littermates. The right panels show tubular injury score (top) and semi-quantitative analysis of the Picro-Sirius red^+ve^ area in the indicated genotypes (bottom) (n=7-8). Scale bars indicate 50 μm. **(c)** *Loxl2*, *Tgfb1*, *Acta2*, and *Havcr1* mRNA levels in IR and CTL kidneys from *PHD1^iEC^* mice and their *Cre −* controls on day 14 after uIRI (n=7). All bars show mean ± SEM. For (**b**), unpaired t-test with Welch’s correction was used. For (**c**), statistics were determined using one-way ANOVA with Sidak correction for multiple comparisons. ns, not statistically significant. uIRI, unilateral ischemia reperfusion injury; CTL, contralateral; IR, kidney subjected to uIRI; Rel., relative.

Next, we generated *Cdh5-CreER^T2^; Phd3 ^f/f^* mice (*PHD3^iEC^*) and subjected them to uIRI, followed by tamoxifen treatment and analysis on day 14 post-uIRI (Figure 3a). Histological analysis of day 14 post-IRI kidneys revealed that compared to *Cre-*littermate controls, *PHD3^iEC^* mutants showed higher tubular injury, as indicated by increased tubular dilation, cast formation, and necrosis (Figure 3b). Moreover, Picro-Sirius red staining indicated increased collagen deposition by ∼42% in post-ischemic *PHD3^iEC^* mutant kidneys compared to *Cre-* littermates (n=6, P <0.05) (Figure 3b). Consistent with the increased post-ischemic kidney fibrosis and injury in *PHD3^iEC^* mutants, significant increases in the mRNA levels of *Loxl2* (P <0.01), *Tgfb1* (*P* <0.01), *Acta2* (*P* <0.05), and *Havcr1*(*P* <0.01) were observed (n=5-6) (Figure 3c). Furthermore, we examined capillary density by endomucin (EMCN) staining and found a 53% reduction in the EMCN^+^ area on day 14 post-uIRI kidneys from *PHD3^iEC^* mice relative to *Cre-* controls (n= 6, *P*<0.01) (Figure 3d). In summary, while post-ischemic inactivation of endothelial PHD1 does not alter kidney repair, post ischemic EC-specific inactivation of PHD3 leads to maladaptive kidney repair, characterized by tubular injury, interstitial fibrosis, and peritubular capillary rarefaction.

**Figure 3.**
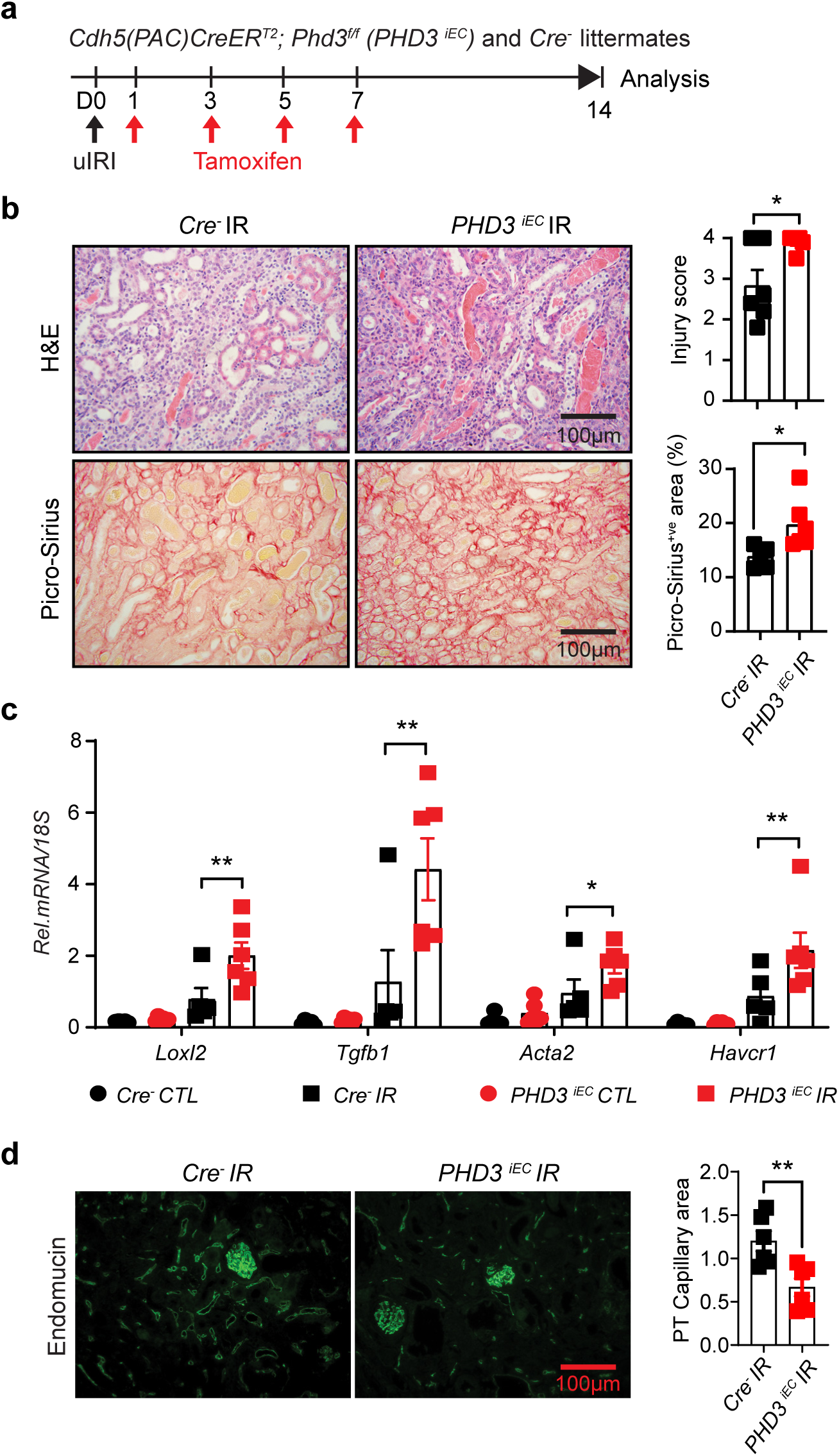
Post-ischemic inactivation of endothelial PHD3 leads to maladaptive kidney repair with increased fibrosis and capillary rarefaction. (**a**) Experimental scheme illustrates the timing of renal artery clamping, tamoxifen administration, and analysis. (**b**) Shown are representative images of H&E- and Picro-Sirius red-stained sections as well as tubular injury score and semi-quantitative analysis of Picro-Sirius red-positive area for day 14 post-IRI kidneys from *PHD3^iEC^*mice and *Cre-* littermates (n=6). Scale bars indicate 50μm. (**c**) mRNA levels of *Loxl2*, *Tgfb1*, *Acta2,* and *Havcr1* in IR and CTL kidneys from *PHD3^iEC^* mice or their *Cre-* controls on day 14 after uIRI (n=5-6). (**d**) Representative images of EMCN immunostaining and semi-quantitative analysis of the EMCN^+ve^ area of day 14 post-uIRI kidneys from *PHD3^iEC^* mice and *Cre-*littermates. Data are represented as mean ± SEM (n=5-6 for each group). For (**b**) and (**d**), statistics were determined by unpaired t-test with Welch’s correction. For (**c**), one-way ANOVA with Sidak correction for multiple comparisons was used. *, *P*< 0.05; **, *P*< 0.01; ns, not statistically significant. CTL, contralateral kidney; IR, kidney subjected to uIRI; Rel., relative.

### Post-ischemic endothelial inactivation of PHD3 induces EC derived pro-inflammatory responses and macrophage infiltration

Because inflammation is emerging as a driver of maladaptive kidney repair, we next sought to investigate whether post-ischemic endothelial PHD3 inactivation alters inflammatory responses. To this end, we first assessed the transcript levels of EC adhesion molecules and other proinflammatory genes in post-ischemic *PHD3^iEC^* mutant and *Cre-* control kidneys. Notably, we found that post ischemic endothelial PHD3 inactivation significantly upregulated intercellular adhesion molecule1 (*Icam1*) (*P* <0.001), vascular cell adhesion molecule1 (*Vcam1*) (*P* <0.01), C-X-C motif chemokine ligand 10 *(Cxcl10)* (*P* <0.05) and macrophage marker *F4/80* (*P* <0.05) transcript levels (n=5-6, Figure 4a). A membrane-based multiplex antibody array was used to confirm the relative levels of selected cytokines in post-ischemic kidney tissues. Consistent with the mRNA levels, the protein levels of ICAM1 and VCAM1 were increased by 2.5- and 3.5- fold, respectively, in the *PHD3^iEC^* post-ischemic kidney compared to the *Cre-* control (Figure 4b). Furthermore, we observed increased levels of other proteins, such as insulin-like growth factor binding protein (IGFBP2), implicated in regulating hypoxia responses during human adaptation to high altitude ^28^ and retinol-binding protein 4 (RBP4), shown to promote vascular inflammation ^29^. Next, we performed flow cytometry analysis on day 14 post-ischemic kidneys from *PHD3^iEC^* and *Cre-* controls. Among the different immune cell populations, macrophages (CD45^+^ CD11b^+^ F4/80^+^) were increased by ∼50% on day 14 post-uIRI *PHD3^iEC^* kidneys relative to *Cre-* controls (n=3-4, *P* <0.05) (Figure 4c). Consistent with the flow cytometry data was the significant upregulation of *F4/80* gene by qPCR in *Phd3^iEC^* mutants compared to *Cre-* littermate controls (n=5-6, *P* <0.05) (Figure 4a). Taken together, these data suggest that post-ischemic endothelial PHD3 inactivation promotes EC-driven inflammatory responses and leads to increased macrophage infiltration in the post-ischemic kidney.

**Figure 4.**
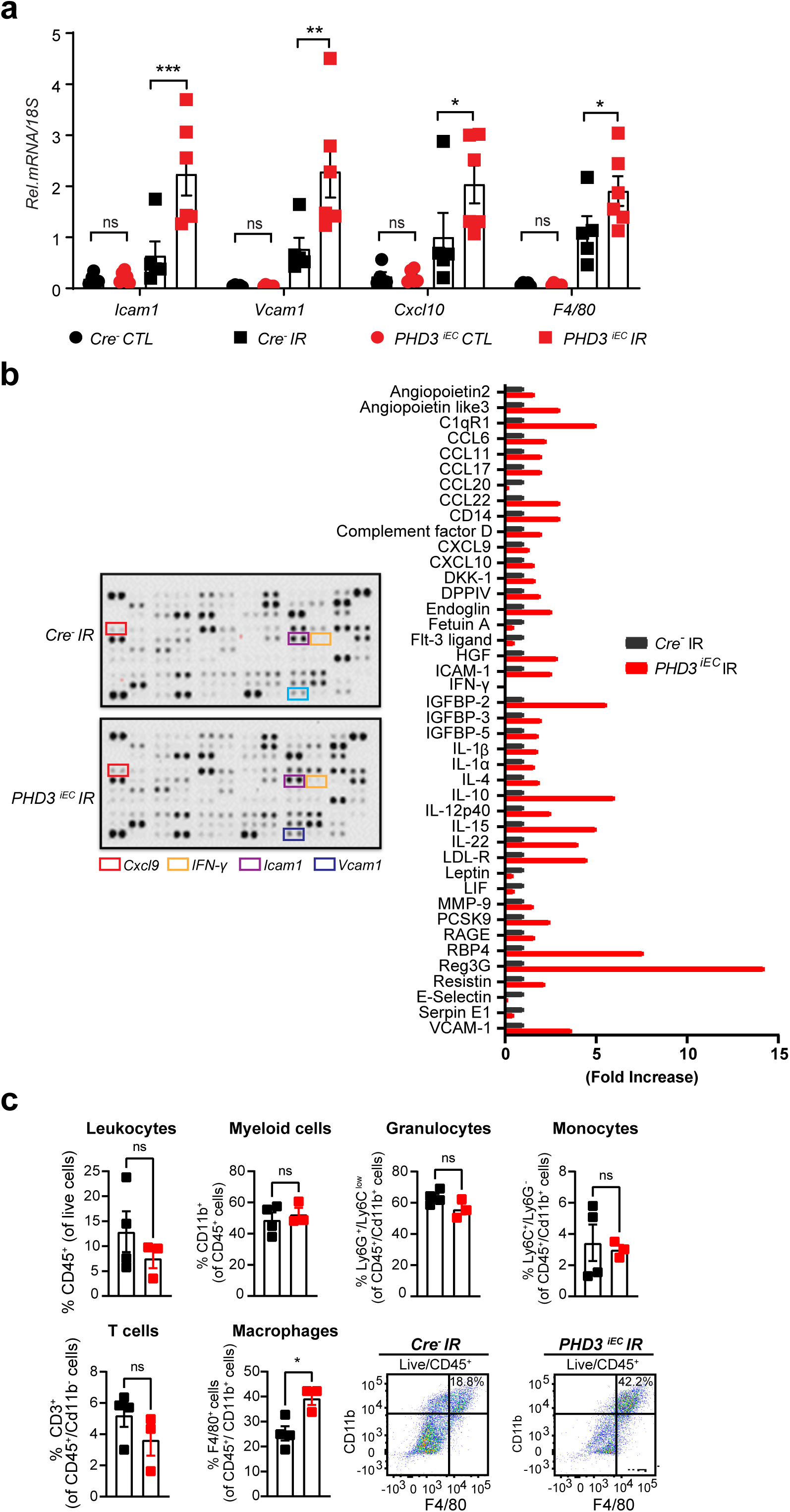
Post-ischemic inactivation of endothelial PHD3 induces endothelial cell-derived pro-inflammatory responses. (**a**) mRNA levels of pro-inflammatory genes *Icam1*, *Vcam1*, *Cxcl10*, and *F4/80* in IR and CTL kidneys from *PHD3^iEC^* mice or their *Cre-* controls at day 14 after uIRI (n=5-6). (**b**) Shown are the representative blots of the antibody cytokine array from *PHD3^iEC^* and *Cre-* kidney tissue homogenates. The bar graph on the right shows the densitometric analysis (n = 1 per genotype). (**c**) Day 14 post-uIRI kidneys from *PHD3^iEC^* and *Cre-* control mice were harvested, and flow cytometry analysis for immune cells was performed (n=3-4). Data are represented as the mean ± SEM. For (**a**) Statistics were determined by one-way ANOVA with Sidak correction for multiple comparisons. For (**c**) unpaired t-test with Welch’s correction was used. *, *P* <0.05; ** *P,*<0.01; ***, *P*< 0.001; ns, not statistically significant. CTL, contralateral kidney; IR, kidney subjected to uIRI; Rel., relative.

### scRNA-seq reveals distinct endothelial cell-specific responses in post-ischemic endothelial PHD3 haplodeficient kidney

To examine how post-ischemic endothelial PHD inactivation influences EC-specific changes during kidney repair , we performed scRNA-seq 14 days post-IRI. We opted to use heterozygous mice to observe early transcriptional changes linked to partial PHD3 deficiency [*Cdh5(PAC)CreER^T2^; Phd3^+/f^* mice, hereafter referred to as *PHD3+/- ^iEC^*]. This strategy enabled us to focus on endothelial-specific alterations while minimizing the confounding effects of severe fibrosis and ensuring high-quality single-cell data. Using the 10X Genomics Chromium platform, we aimed for 10,000 cells per group. After quality filtering, 19,441 (9,440 from *Cre^-^* and 10,001 from *PHD3+/- ^iEC^*) cells were retained for subsequent analysis. We identified 11 different cell clusters using unsupervised graph-based clustering followed by principal component analysis (PCA) and uniform manifold approximation and projection (UMAP) (Figure 5a). When we overlaid the *Cre^-^*IR and *PHD3+/- ^iEC^* IR samples, we observed similar cell clusters for each genotype (Supplemental Figure 3). Based on expression of reported canonical marker genes ^20^, we identified the following kidney cell types; proximal tubule (PT), loop of Henle (LOH), collecting duct 1 and 2 (CD1 and CD2), endothelial cells (EC), parietal cells (PAR), fibroblasts/smooth muscle cells (FIB/SMC), macrophages 1 and 2 (Mφ1 and Mφ2), neutrophils (NEU), and T-cells (T) (Figure 5b). Next, we performed differential gene expression analysis for the EC cluster between the two genotypes. After applying cutoff criteria of a log_2_ fold change >0.2 and adjusted *P*<0.05, EC showed 278 differentially expressed genes (DEGs) (Supplemental Data S1). Gene set enrichment analysis (GSEA) of DEGs demonstrated significant differences in EC between *PHD3+/- ^iEC^* and *Cre^-^*. Notably, the top Hallmark gene signatures included Interferon Gamma Response, Myc Target V1, and Interferon Apha Response (Figure 5c). Consistent with the enhanced Interferon Gamma Response was the significant upregulation of genes such as interferon-induced transmembrane protein (*Ifitm3*), interferon-induced protein with tetratricopeptide repeats 3 (*Ifit3*), and Interferon regulatory factor 7 (*Irf7*) in EC from *PHD3+/-^iEC^*IR kidney compared to *Cre^-^* control (Figure 5d). Overall, these results demonstrate that post-ischemic endothelial inactivation of PHD3 activates pro-inflammatory responses with a prominent IFN-γ signature.

**Figure 5.**
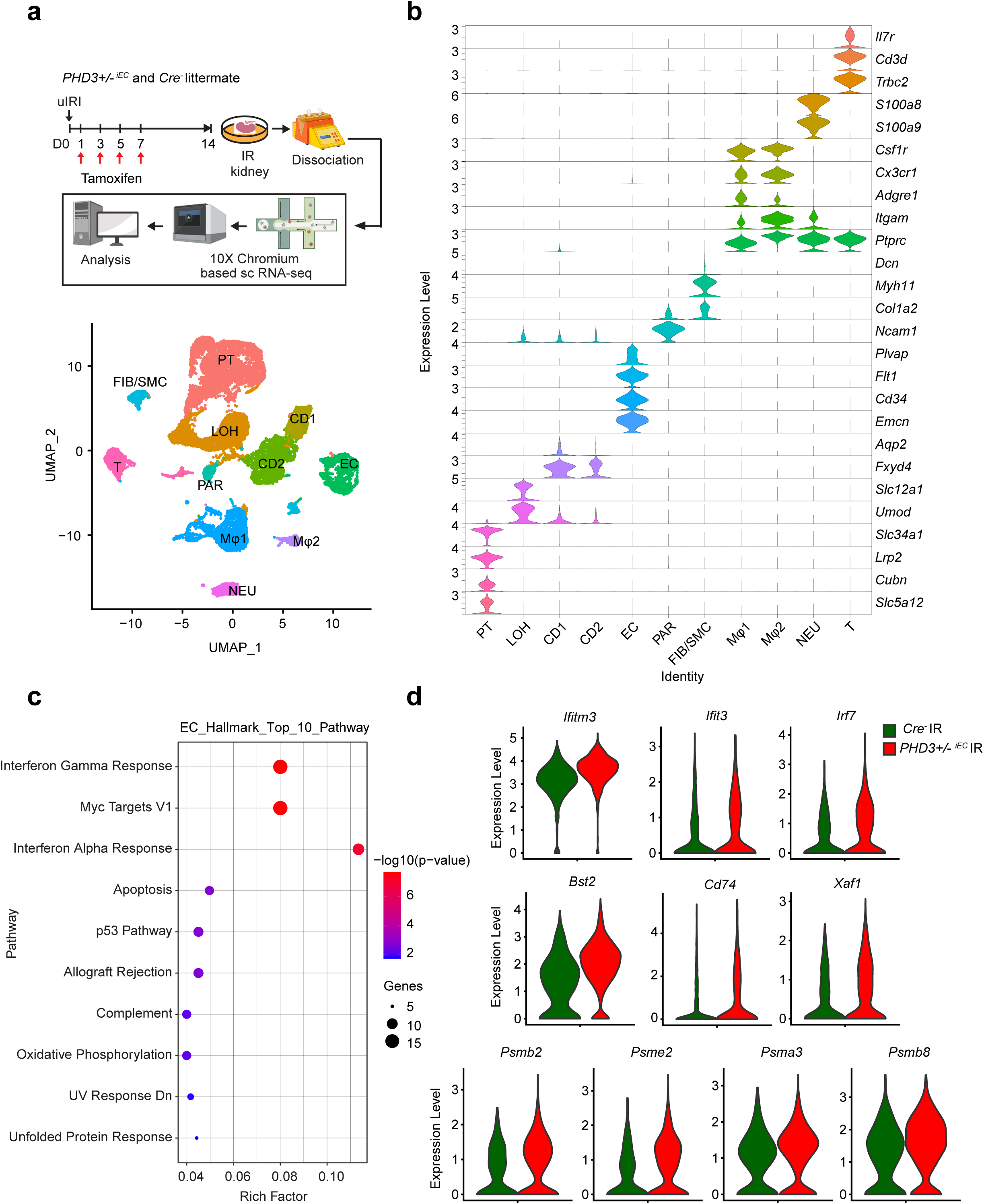
scRNA-seq analysis reveals the activation of endothelial Interferon-γ response on day 14 post-ischemic kidney in the setting of endothelial PHD3 haplodeficiency. **(a)** Top: Scheme illustrating the experimental strategy applied for scRNA-seq analysis. A *PHD3+/- ^iEC^* mouse and a *Cre^-^* littermate were subjected to 25 min of unilateral renal artery clamping. Treatment with tamoxifen was started on day 1 post uIRI and involved 4 i.p. doses administered every other day. Mice were sacrificed for scRNA-seq analysis on day 14 post-uIRI. IR kidneys were isolated, single-cell suspensions were prepared, and used for scRNA-seq analysis (n=1 per genotype). Bottom: Uniform manifold approximation and projection (UMAP) plot representation of cell classification on day 14 post-ischemic kidneys from *PHD3+/- ^iEC^* and *Cre^-^* mice. **(b)** Violin plots display characteristic marker genes for each identified cell population. Proximal tubule (PT), Loop of Henle (LOH), Collecting duct 1 and 2 (CD1 and CD2), Endothelial cells (EC), Parietal cells (PAR), Fibroblasts/Smooth muscle cells (FIB/SMC), Macrophages 1 and 2 (Mφ1 and Mφ2), Neutrophils (NEU) and T cells (T). **(c)** Top 10 enriched Hallmark pathways emerged in Hallmark gene set enrichment analysis (GSEA) of differentially expressed genes (DEGs) for the EC cluster of *PHD3+/- ^iEC^* post-ischemic kidney as compared to *Cre^-^* control. **(d)** Violin plots show significantly upregulated Interferon Gamma Response genes in the EC cluster of *PHD3+/- ^iEC^* compared to the control.

### Endothelial PHD3 inhibition in primary endothelial cells regulates IFN-γ induced pro-inflammatory transcriptional responses

To assess whether PHD3 inhibition acts cell-autonomously in EC to regulate pro-inflammatory responses in a HIF-dependent manner, we performed in vitro experiments in which human primary pulmonary artery EC (HPAECs) were treated with PHD3 siRNA alone or double transfected with PHD3 and ARNT siRNA for 48 h (Figure 6a). With more than ∼50% knockdown efficiency for both siRNAs, we observed that siPHD3 treatment led to transcriptional induction of *ICAM1* (*P* <0.01), *VCAM1* (*P* <0.001), and other pro-inflammatory genes in ECs compared to negative control siRNA, whereas simultaneous ARNT inactivation blunted the effect of PHD3 inhibition (Figure 6b).

**Figure 6.**
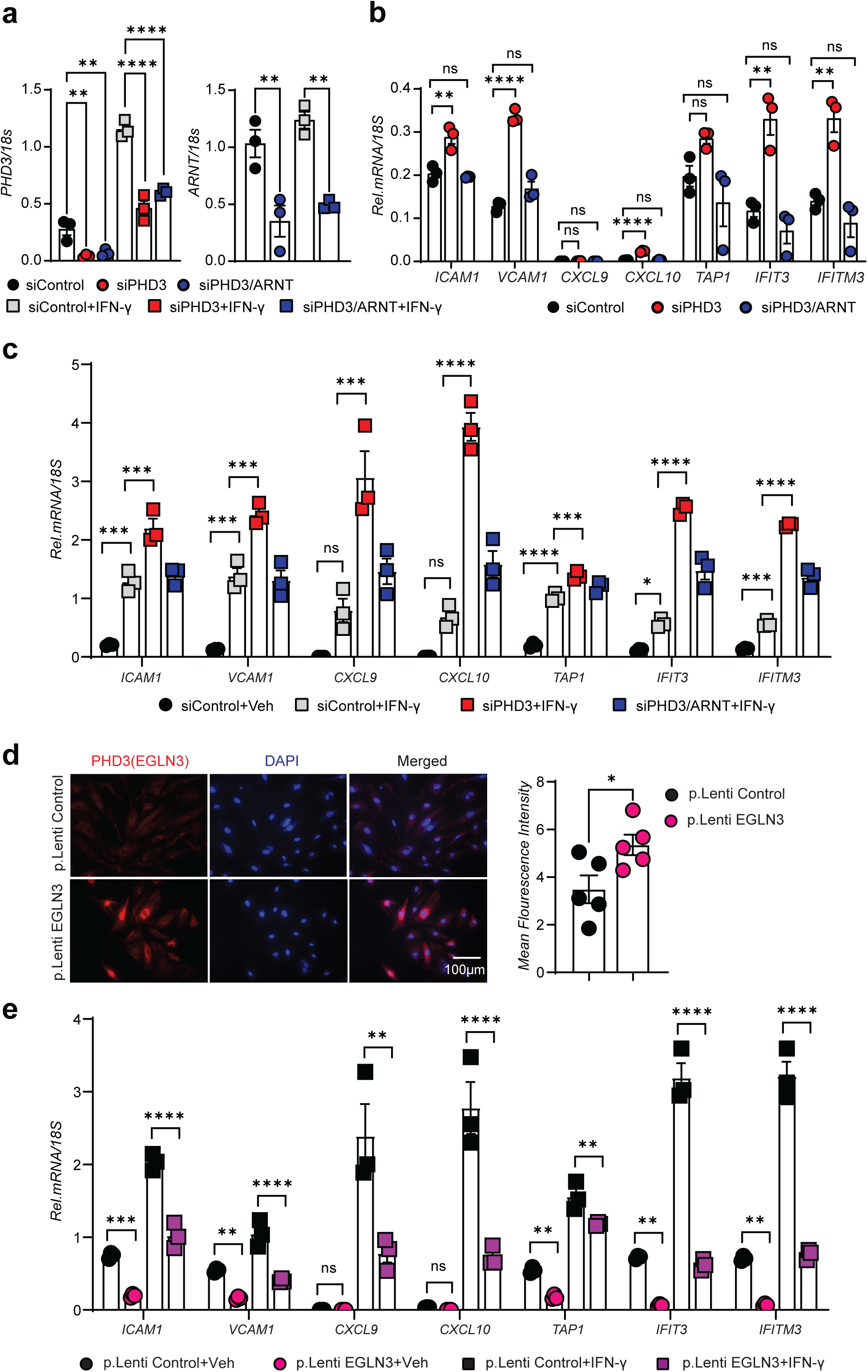
PHD3 regulates the transcription of IFN-γ-responsive pro-inflammatory genes in an ARNT-dependent manner. (**a**) Gene expression analysis by RT-PCR was performed at baseline or after IFN-γ stimulation. Shown are *PHD3* and *ARNT* mRNA levels in cells transfected with siRNAs against *PHD3* and *ARNT* compared to negative control siRNA-transfected cells. (**b**) mRNA expression levels of pro-inflammatory genes *ICAM1*, *VCAM1*, *CXCL9*, *CXCL10*, *TAP1*, *IFIT3*, and *IFITM3* in HPAECs transfected with control (scrambled), si*PHD3*, or double transfected with si*PHD3* and si*ARNT* for 48 h at baseline conditions and following IFN-γ stimulation (**c**). (**d**) Representative images of immunofluorescence staining for PHD3 (red) and nuclear DAPI (blue) in HPAECs transduced with *p.Lenti.EGLN3* or *p.Lenti.Control.* Scale bar, 100µm. The graph on the left panel shows the mean florescence intensity quantitation of EGLN3 expression in lentivirus-transduced HPAECs. (**e**) mRNA expression levels of pro-inflammatory genes *ICAM1*, *VCAM1*, *CXCL9*, *CXCL10*, *TAP1*, *IFIT3*, and *IFITM3* in HPAECs transduced with *p.Lenti.EGLN3* or *p.Lenti.Control* at baseline or after IFN-γ stimulation (n=3). All bars show mean ± SEM. For (**a**), (**b**), (**c**), and (**e**), statistics were determined by one-way ANOVA with Sidak correction for multiple comparisons. For (**d**), unpaired t-test with Welch’s correction was used. *, *P*<0.05 ; **, *P*<0.01; ***, *P*<0.001 ; ****, *P*< 0.0001; ns, not statistically significant.

We next focused on the possibility that PHD3 may play an important role in regulating responses to pro-inflammatory stressors. Because IFN-γ induces PHD3 ^30^ and IFN-γ responses were enriched in *PHD3* haplodeficient EC after IRI, we hypothesized that PHD3 could be a crucial molecular link between hypoxia and IFN-γ-signaling. We first confirmed that *PHD3* mRNA was upregulated in EC stimulated with IFN-γ (Supplemental Figure 4), as previously reported ^30^. To investigate whether PHD3 regulates the endothelial transcriptional response to IFN-γ, we used a siRNA approach to inhibit PHD3 inhibition in the presence of IFN-γ (Figure 6c). We found that PHD3 inhibition significantly boosted IFN-γ-induced expression of *ICAM1* (∼1.7-fold), *VCAM1* (∼1.7-fold), *CXCL9* (∼3.8-fold), *CXCL10* (∼5.7-fold), *IFIT3* (∼4.4-fold), *IFITM3* (∼3.9-fold), and *TAP1*(∼1.3-fold) (Figure 6c). Notably, the effects of PHD3 inhibition were abrogated when ARNT was concurrently inhibited (Figure 6c). Therefore, inhibition of PHD3 exacerbates the endothelial transcriptional response to IFN-γ in an ARNT-dependent manner.

To further investigate the role of PHD3 on IFN-γ-induced EC responses, we performed gain-of-function in vitro studies. Specifically, we used a lentiviral-based approach to transduce HPAECs with pLp6.3. EGLN3, while control cells were transduced with pLp6.3. Control, which encodes a reverse EGLN3 insert (Figure 6D). Immunofluorescence staining showed a ∼44% increase in PHD3 protein expression levels in EGLN3-transduced HPAECs relative to cells transduced with the negative control plasmid (Figure 6d). Importantly, overexpression of PHD3 in HPAECs significantly abrogated the upregulation of EC cell adhesion molecules, *ICAM1* (p <0.0001), *VCAM1* (p <0.0001), and other IFN-γ responsive genes such as *CXCL9* (*P* <0.01), *CXCL10* (*P* <0.0001), *IFIT3* (*P* <0.0001), *IFITM3* (*P* <0.0001), and transporter associated with antigen processing 1 (*TAP1*) (p<0.01) (Figure 6e). Since activation of HIF was shown to induce the transcription of apolipoprotein 1 (*APOL1*) ^31^, we next assessed whether PHD3 regulates *APOL1* in EC. PHD3 inhibition significantly upregulated endothelial *APOL1* transcript and enhanced the effect of IFN-γ on inducing *APOL1* in an *ARNT*-dependent manner (Supplemental Figure 5a). In contrast PHD3 overexpression suppressed *APOL1* mRNA in this setting (Supplemental Figure 5b). Therefore, these results identify PHD3 as a critical regulator of the endothelial pro-inflammatory responses to IFN-γ.

### Post ischemic concurrent inactivation of endothelial PHD3 and ARNT partially rescues the pro-fibrotic phenotype mediated by inactivation of endothelial PHD3

To assess the potential role of HIF signaling in the context of endothelial PHD3 inhibition, we first assessed HIF-1 and HIF-2 stabilization in the kidney tissue of *PHD3^iEC^* mice at baseline conditions and following IRI. Nuclear extracts from day 14 post-IRI kidneys showed a significant increase in HIF-1α protein levels in mutant *PHD3^iEC^* kidneys compared to *Cre-* controls, while in the absence of injury, there was no difference between the two genotypes (Supplemental Figure 6a&b). Next, to test whether HIF signaling is required in the maladaptive kidney repair induced by inhibition of endothelial PHD3, we used a genetic approach in which HIF signaling is ablated by deletion of *ARNT* in the context of endothelial PHD3 inhibition (Figure 7a). Specifically, we generated *PHD3ARNT^iEC^* double mutant mice by crossing *Cdh5(PAC)-CreER^T2^;PHD3^f/f^* (*PHD3^iEC^*) mice with *ARNT ^f/f^* mice. Notably, *PHD3ARNΤ^iEC^* kidneys had similar tubular damage to *Cre-* littermate controls (Figure 7b). Consistently, quantitative analysis of Picro-Sirius red staining showed no significant difference in collagen deposition between *PHD3ARNT ^iEC^* and *Cre-* controls (Figure 7b). Furthermore, there was no significant difference in the transcript levels of *Loxl2, Tgfb1*, and *Havcr1* between *PHD3ARNT ^iEC^* and controls, while *Acta2* remained significantly increased in *PHD3ARNT ^iEC^*mutants (n=5-6, ns) (Figure 7c). Furthermore, the expression of pro-inflammatory genes *Vcam1*, *Icam1*, *Cxcl10*, and *F4/80* was comparable between *PHD3ARNT ^iEC^* and controls (Figure 7d), in contrast to the increases noted in the single *PHD3^iEC^* mutants (Figure 4a). Therefore, by generating mice with combined endothelial deletion of PHD3 and ARNT, we demonstrated that the pro-fibrotic effect of endothelial PHD3 inhibition in postischemic AKI depends on HIF signaling.

**Figure 7.**
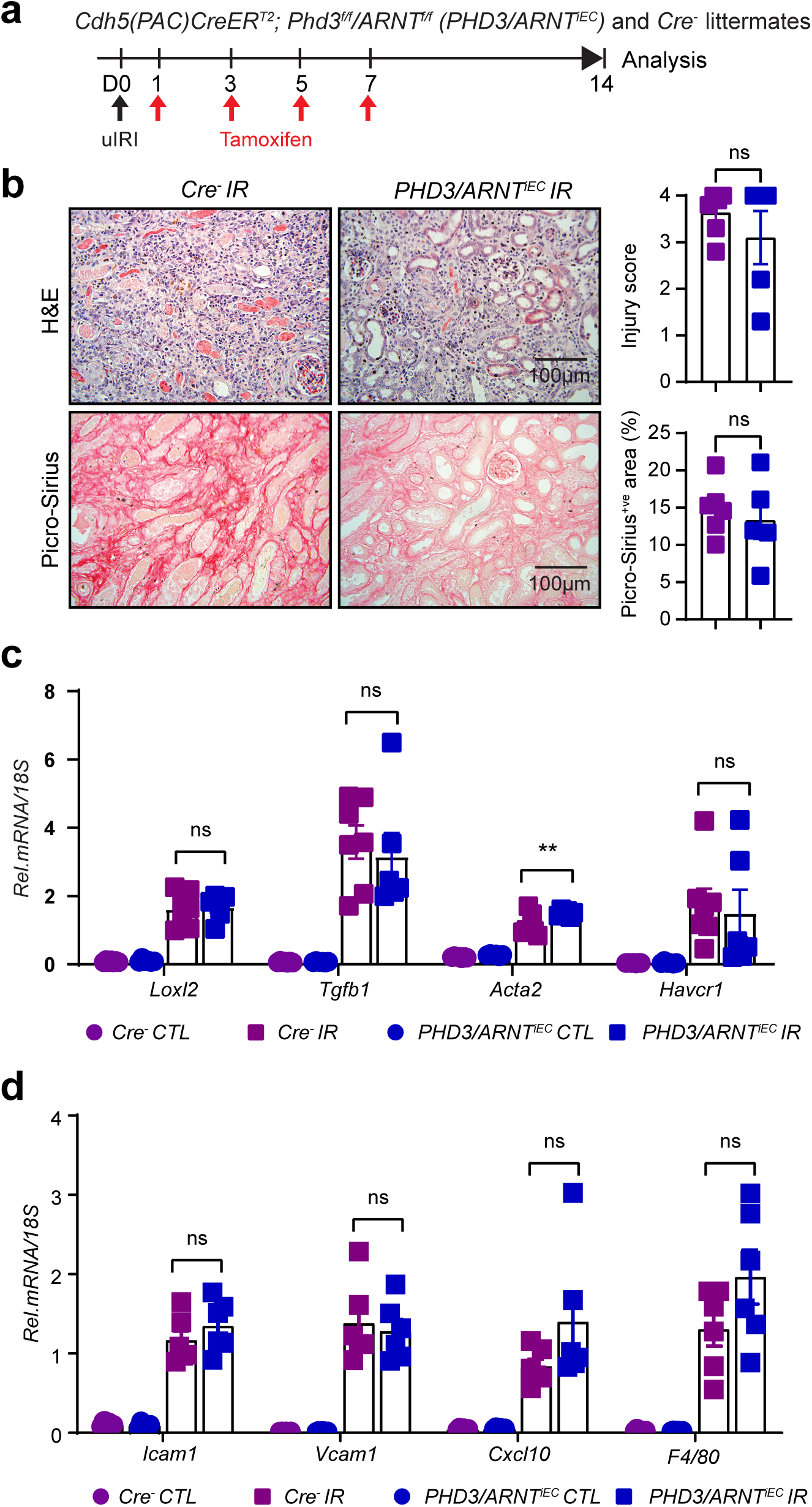
Post-ischemic inactivation of endothelial ARNT blunts the pro-fibrotic effect induced by endothelial PHD3 loss following kidney IRI. (**a**) Schematic depicting the experimental approach used. (**b**) Representative images of H&E- and Picro-Sirius red-stained sections as well as tubular injury score and semi-quantitative analysis of Picro-Sirius red^+ve^ area on day 14 post-IRI kidneys from *PHD3ARNT ^iEC^* mice and *Cre-* littermates. Scale bars indicate 100 μm and 200 μm for H&E and Picro-Sirius red images, respectively. mRNA levels of profibrotic genes *Loxl2*, *Tgfb1*, *Acta2,* and *Havcr1* (**c**) and pro-inflammatory genes *Icam1*, *Vcam1*, *Cxcl10, and F4/80* (**d**) in IR and CTL kidneys from *PHD3ARNT ^iEC^* mice or their *Cre-* controls at day 14 post uIRI. All bars show mean ± SEM (n=6-7 for each group). For (**b**), statistics were determined by unpaired t- test with Welch’s correction. For (**c**) and (**d**), one-way ANOVA with Sidak correction for multiple comparisons was used.**, *P*< 0.01; ns, not statistically significant. CTL, contralateral kidney; IR, kidney subjected to uIRI; Rel., relative.

Finally, we assessed whether the regulation of EC responses by PHD3/HIF signaling is relevant to human AKI. To this end, we analyzed publicly available snRNA-seq data from kidney tissues of patients with severe AKI ^26^. IPA analysis of DEGs in kidney EC from AKI patients compared to controls showed activation of HIF1 signaling and IFN-γ as key molecular responses of their dysregulated state (Supplemental Figure 7). Overall, this analysis demonstrates the relevance of endothelial IFN-γ signaling and hypoxia signaling in human AKI.

## DISCUSSION

Here, using a genetic approach in mice subjected to a model of AKI-to-CKD transition, we sought to investigate how alterations in endothelial oxygen sensing regulate the kidney microenvironment and shape repair outcomes. We found that post-ischemic inactivation of endothelial PHD3, but not PHD1, leads to maladaptive kidney repair characterized by exacerbated injury, fibrosis, and inflammation. Guided by previous studies on hypoxia biology and kidney repair, we examined the molecular changes that might potentially underlie the effects of PHD3 in the kidney endothelium. Our mechanistic studies revealed that endothelial PHD3 determines postischemic kidney repair by a HIF-dependent mechanism that regulates IFN-γ-driven pro-inflammatory signaling.

PHDs are critical oxygen sensors; however, their roles in kidney homoeostasis and kidney disease remain incompletely understood. Combined inactivation of stromal PHD2 and PHD3 resulted in renal failure, associated with decreased numbers of glomeruli and abnormal postnatal nephron formation, findings that underscore the profound impact of PHDs in kidney development^32^. Furthermore, PHD2 and PHD3 were shown to function as the main regulators of renal erythropoietin (EPO)-producing cell plasticity, dictating systemic EPO levels ^33^. Combined inducible endothelial deletion of PHD1, PHD2, and PHD3 worsened kidney injury and fibrosis without affecting baseline kidney homeostasis, whereas post-ischemic PHD2 deletion alone had no impact on repair ^20^. Here, we identified endothelial PHD3 as a crucial regulator of post-ischemic kidney repair, whereas endothelial PHD1 was dispensable. Consistently, PHD3 deficiency aggravated the innate immune responses to sepsis ^34^. Nevertheless, given the renoprotective effects by pharmacological pan-PHD inhibitors given prior to AKI, it is likely that compensatory effects from other cell types or PHD isoforms counterbalance the detrimental impact of endothelial-specific PHD3 loss. In support of a cell-type-specific and context-dependent role for PHD3 in dictating injury outcomes are studies from other organs and tumor biology. For instance, systemic deletion of PHD3 before myocardial IRI exerted protection by inhibiting the DNA damage response, leading to decreased cardiomyocyte apoptosis ^35^. Additionally, PHD3 promotes apoptosis in neurons and other cell types under normoxic conditions ^36^, whereas under hypoxia, it supports neutrophil survival ^37^ and enhances cell cycle progression in cancer cells ^38^.

Although multiple inputs regulate HIF, hydroxylation primarily controls its degradation and activity, and therefore PHD isoform levels dictate HIF activation. By investigating the expression of PHD isoforms in murine and human AKI, we found increased expression of PHD3 in kidney EC. This response keeps up with the well-established hypoxic regulation of PHD3 by HIF ^19^, but could also be driven independently of HIF through cytokines such as TNF-α and IL-1β^39^. What is the biological significance of increased endothelial PHD3 abundance in AKI? Elevated PHD3 levels enable EC to fine-tune the threshold for HIF activation, thereby enhancing their ability to regulate and adapt to reduced oxygen availability ^40^. Indeed, we found that lack of endothelial PHD3 led to increased HIF stabilization in the post-ischemic kidney. Furthermore, inactivating HIF in the context of endothelial PHD3 deficiency improved kidney repair, supporting a role for PHD3 in restraining maladaptive hypoxic signaling. Since excessive endothelial glycolysis due to aberrant HIF activation is a driver of maladaptive repair ^20^, PHD3 may suppress endothelial glycolysis by HIF hydroxylation. Nevertheless, non-canonical mechanisms may also be involved. For instance, PHD3 has been shown to suppress glycolysis by direct binding to pyruvate kinase ^41^, which could generate a more immediate response aiming to adjust metabolism according to oxygen availability.

The injured kidney tissue microenvironment is shaped by hypoxia-inflammation crosstalk, with cytokines like IFN-γ driving immune responses and influencing disease progression through chemokine induction and endothelial dysfunction. Gerber et al. demonstrated endothelial PHD3 induction by IFN-γ and suggested that PHD3 enhances IFN-γ signaling. This assumption was based on the suppression of IFN-γ transcriptional responses by two non-selective PHD inhibitors, dimethyloxalylglycine and deferoxamine ^30^. While they sought to further investigate whether the effects of PHD inhibitors were specifically due to PHD3 inhibition, they were limited by insufficient knockdown of PHD3 in HUVECs. Therefore, our studies provide crucial clarity on this unresolved question, demonstrating that PHD3 functions as a key inhibitor of IFN-γ signaling via a HIF-dependent mechanism. Intriguingly, IFN-γ suppresses HIF-1 activity by repressing ARNT directly in intestinal epithelial cells ^42^ and indirectly in EC through kynurenine-mediated AHR activation. ^43^.

Stabilization of HIF by hypoxia or PHD inhibitors potentiates IFN-γ transcriptional induction of *APOL1* in kidney cells ^31^, a process we now demonstrate is PHD3-dependent in EC. These findings carry important translational implications, particularly for anemic patients harboring APOL1 risk variants treated with PHD inhibitors. Patients carrying APOL1 risk variants are more susceptible to sepsis ^44^, and our findings suggest that non-selective PHD inhibitor therapy could further worsen sepsis outcomes. Furthermore, non-selective PHD inhibition may affect CKD progression by mechanisms that involve IFN-γ-induced endothelial injury and pyroptosis, as recently reported ^45^. Nevertheless, while our genetic model of endothelial PHD3 inactivation offers valuable specificity, it may not recapitulate the pharmacokinetics or selectivity of clinical PHD inhibitors, which inhibit different PHD isoforms in a partial and intermittent manner.

Among the three well-studied proline hydroxylases, PHD3 primarily regulates the target proteins through hydroxylation. In addition to its established role in modifying HIF-α, PHD3 has been shown to hydroxylate other proteins, including p53, acetyl-CoA carboxylase 2 (ACC2), and TFAM, thereby modulating their functions ^46–48^. Notably, PHD3 also influences its interaction partners through mechanisms that are independent of its hydroxylase activity. For instance, PHD3 can stabilize p53 by inhibiting its interaction with MDM2 in a hydroxylation-independent manner^49^. Consistently, PHD3 suppresses *irf7* activity in zebrafish without requiring hydroxylation, but rather diminishes its K63-linked ubiquitination ^50^. Therefore, while our data suggest that endothelial PHD3 deficiency drives maladaptive kidney repair through HIF, we cannot exclude contributions from other substrates or hydroxylation-independent mechanisms.

In summary, our study identifies endothelial PHD3 as a gatekeeper of post-ischemic kidney repair. Through modulation of HIF-dependent mechanism that tempers IFN-γ driven signaling, endothelial PHD3 mitigates maladaptive pro-inflammatory responses and promotes kidney repair. Future studies should focus on refining selective PHD-targeting approaches that promote repair, while minimizing adverse immune consequences.

## DISCLOSURE

The authors declare that they have no conflicts of interest.

## DATA STATEMENT

Mouse scRNAseq data were deposited in the NCBI Gene Expression Omnibus database (GSE GSE294712).

## ACKNOWLEDGMENTS

This work was supported by National Institutes of Health (NIH) grants R01DK115850 and R01DK132672 (P.P.K.) and an American Heart Association (AHA) post-doctoral fellowship 23POST1020467 (R.T.). We acknowledge the Northwestern University George M. O’Brien Kidney Research Core Center (NU GoKidney), an NIH/NIDDK funded program (P30 DK114857), for their core services and support. Imaging work was performed at the Northwestern University Center for Advanced Microscopy, generously supported by NCI CCSG P30 CA060553 awarded to the Robert H Lurie Comprehensive Cancer Center. Sc-RNA-seq was performed at the Northwestern University NUSeq Core Facility. Histology services were provided by the Northwestern University Research Histology and Phenotyping Laboratory, which is supported by NCI P30-CA060553 awarded to the Robert H Lurie Comprehensive Cancer Center. The funders had no role in the study design, data collection and interpretation, or the decision to submit the work for publication.

