## Supplement for "Endothelial Prolyl Hydroxylase 3 Mitigates Maladaptive Inflammation to Promote Post-Ischemic Kidney Repair"

**SUPPLEMENTARY FIGURE LEGENDS**

**Supplementary Figure 1. Differential expression of *PHD* isoforms in kidney ECs in human AKI.** Violin plots show the expression of *PHD2* and *PHD3* genes in kidney ECs in controls vs. AKI patients. The analysis was performed using publicly available snRNA-seq data from Christian Hinze et al [1].

**Supplementary Figure 2. Ischemic kidney injury differentially alters the expression of PHD isoforms in the murine renal endothelium.** Shown are individual channels and merged images from uninjured and post-IRI kidney sections of *Cdh5-Cre;mT/mG* reporter mice at the indicated time points. Red, non-recombined cells expressing membrane bound mTomato; green, recombined cells expressing membrane-bound EGFP; magenta, cells expressing the indicated PHD isoform. Merged images shown here are also presented in Figure 1b. Images were captured using a Nikon Ti2 Widefield fluorescence microscope. Scale bar indicates 50 µm.

**Supplemental Figure 3. scRNA-seq analysis shows similar cell populations on day 14 post-ischemic kidneys of *PHD3+/-* $^{iEC}$ and *Cre*$^{-}$ mice. (a)** UMAP showing the different cell clusters on day 14 post-ischemic kidneys from *PHD3+/-* $^{iEC}$ and *Cre*$^{-}$ mice. **(b)** UMAP after overlaying samples of *Cre*$^{-}$ and *PHD3+/-* $^{iEC}$ day 14 post-ischemic kidneys showing similar clustering.

**Supplementary Figure 4. IFN-γ treatment induces *PHD3*, but not *PHD1* and *PHD2* mRNA levels in HPAECs.** mRNA expression levels of *PHD1, PHD2,* and *PHD3* genes in HPAECs after IFN-γ stimulation. Data are represented as the mean ± SEM. Statistics were determined by unpaired t-test with Welch's correction. *, $P< 0.05$; **, $P< 0.01$ ; ns, not statistically significant.

**Supplementary Figure 5. The PHD3/ARNT signaling axis regulates IFN-γ-dependent induction of *APOL1* in human endothelial cells. (a)** mRNA expression levels of *APOL1* mRNA in HPAECs transfected with control (scrambled), si*PHD3*, or double transfected with si*PHD3* and si*ARNT* for 48 h at baseline conditions and following IFN-γ stimulation. (**b**) *APOL1* mRNA levels in HPAECs transduced with *p.Lenti.EGLN3* or *p.Lenti.Control* at baseline or with IFN-γ stimulation. All bars show mean ± SEM. Statistics were determined by one-way ANOVA with Sidak correction for multiple comparisons. *, $P$<0.05 ; **, $P$<0.01; ***, $P$<0.001 ; ****, $P$< 0.0001; ns, not statistically significant.

**Supplementary Figure 6. Post-ischemic inactivation of endothelial PHD3 stabilizes HIF in mouse post-IRI kidneys**. Immunoblot analysis of HIF-1α and HIF-2α in kidney nuclear extracts isolated from **(a)** $Phd3^{iEC}$ mice under baseline conditions and **(b)** $Phd3^{iEC}$ mice subjected to uIRI and their corresponding *Cre-* littermates. The graphs on the right show the densitometric analysis of normalized HIF protein levels. Ponceau staining (**a**) and detection of histone 3 (**b**) were used for normalization of protein loading. +veC indicates the positive control. Data are represented as the mean ± SEM. Statistics were determined by unpaired t- test with Welch's correction. ∗, $P$ < 0.05; ns, not statistically significant.

**Supplementary Figure 7. Unbiased analysis identified HIF and IFN-γ signaling as key elements of kidney endothelial cell responses in human AKI.** DEGs of the kidney EC cluster of AKI patients that emerged in sn-RNAseq analysis were used to perform the core analysis in QIAGEN Ingenuity Pathway Analysis (IPA) software. The resulting graphical summary network shows major biological changes and their associations. The predicted activity is represented by colored nodes and lines. Orange indicates predicted activation, whereas blue indicates predicted inhibition. Solid lines indicate direct interaction, dashed lines indicate indirect interaction, and dotted lines indicate inferred relationships.

**Supplementry Figure 1**

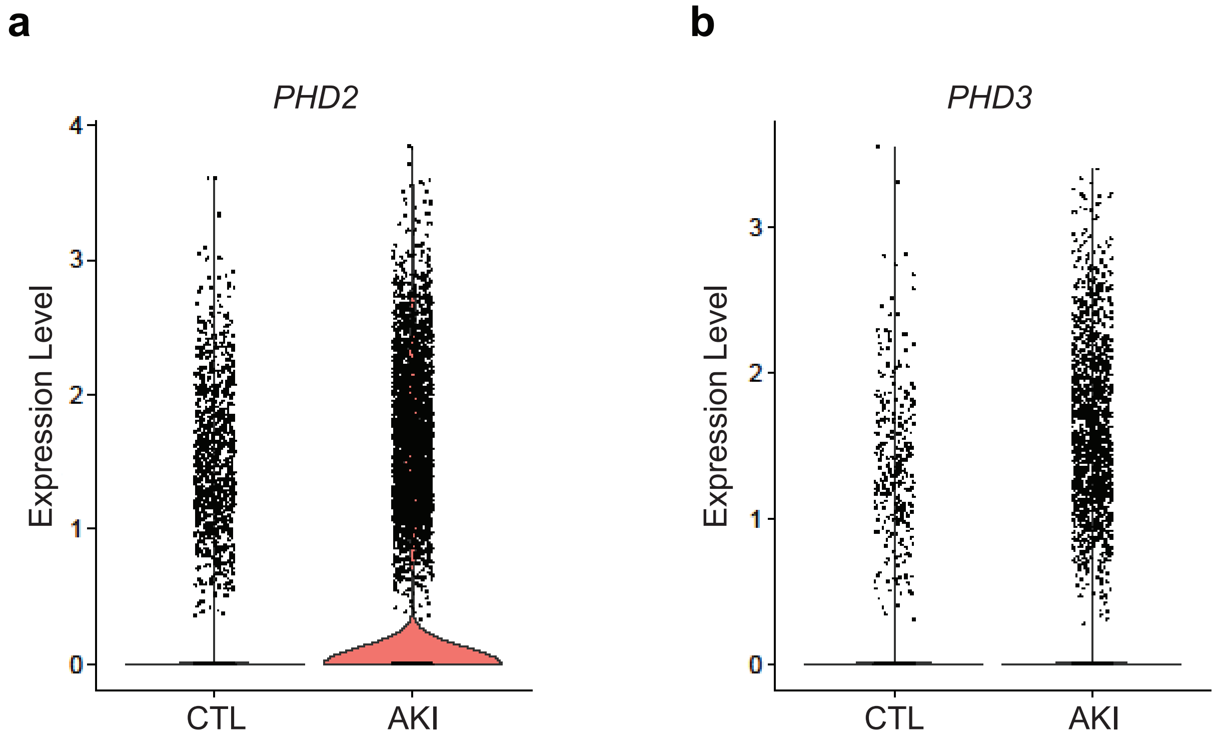

### Supplementry Figure 2

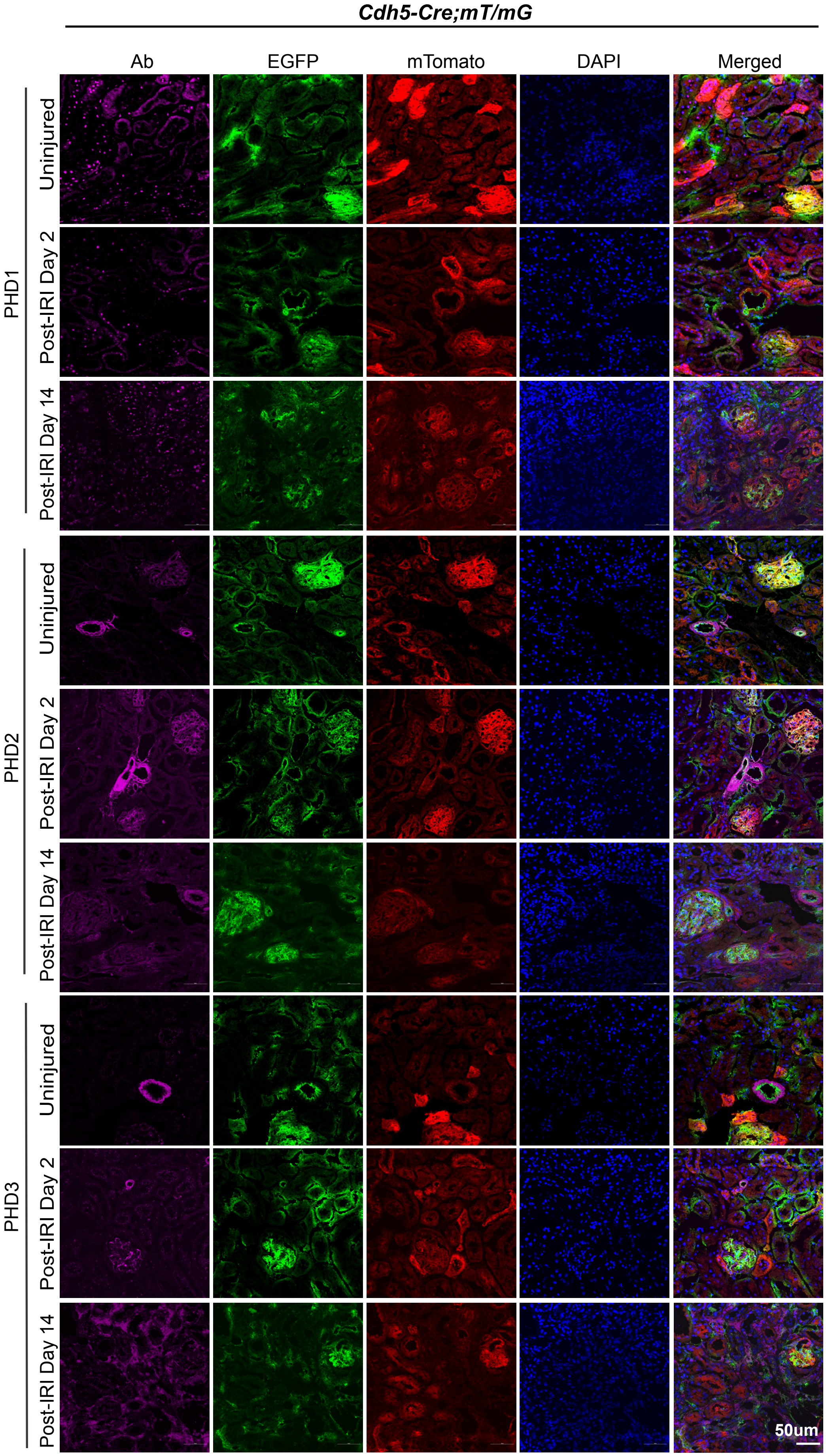

### Supplementry Figure 3

**a**

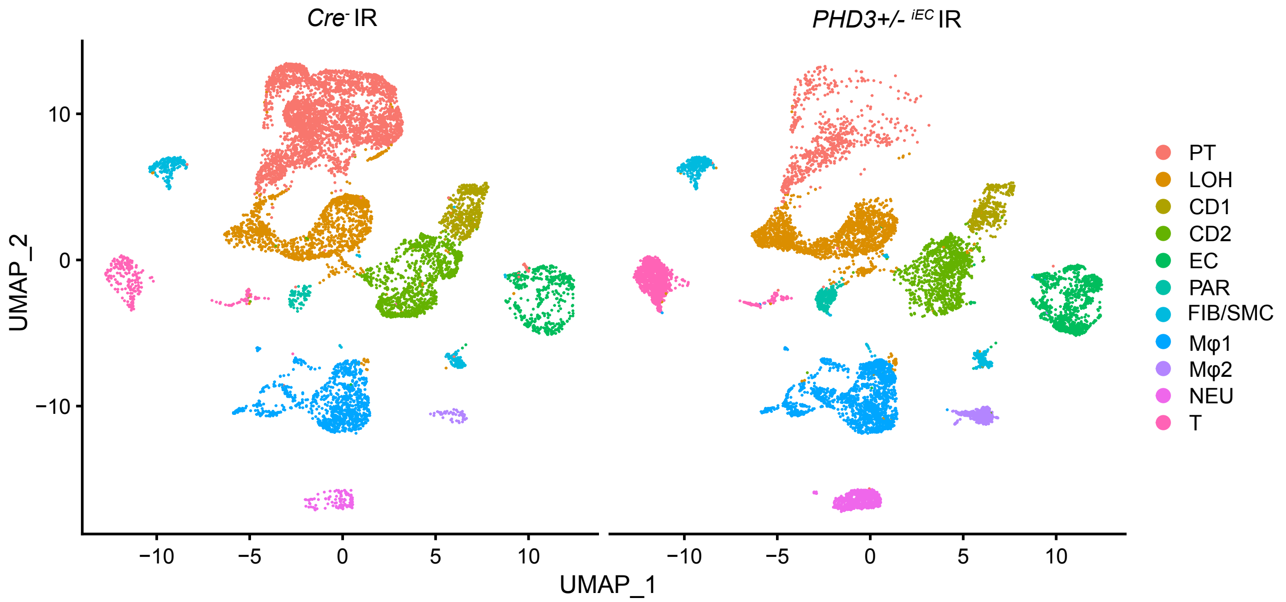


**b**

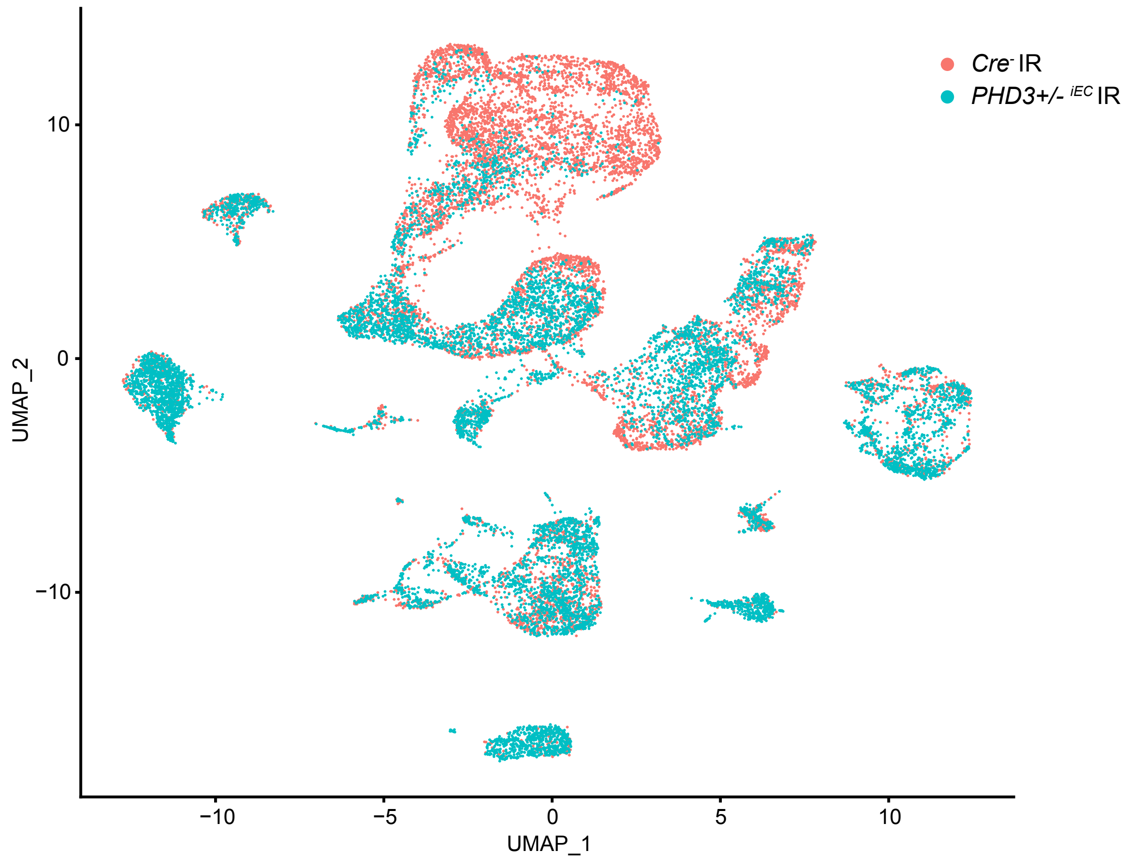

### Supplementry Figure 4

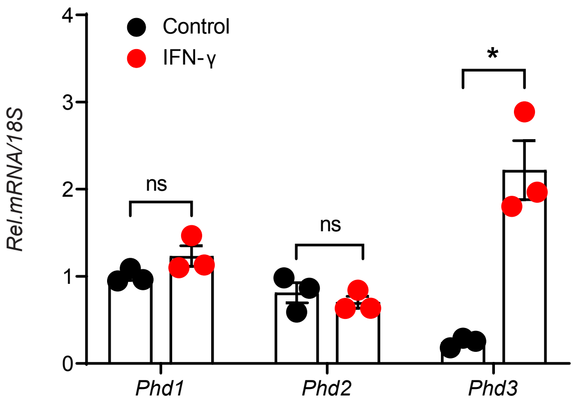

**Supplementry Figure 5**

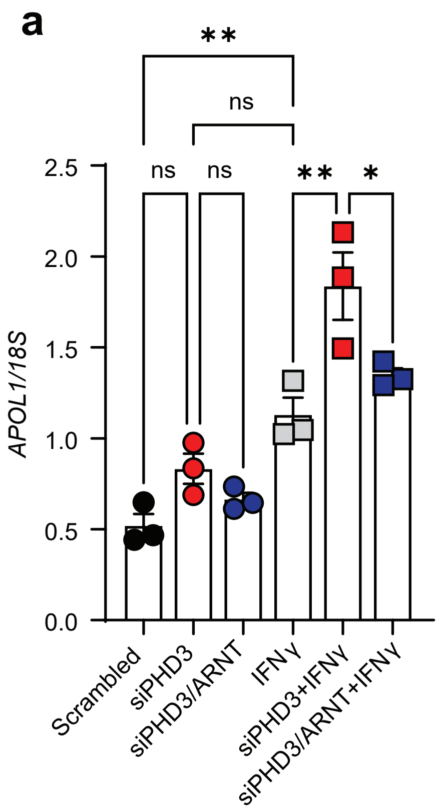


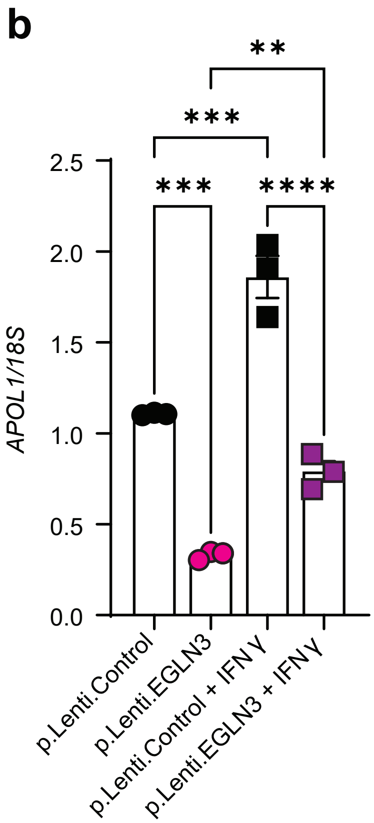

**Supplementry Figure 6**

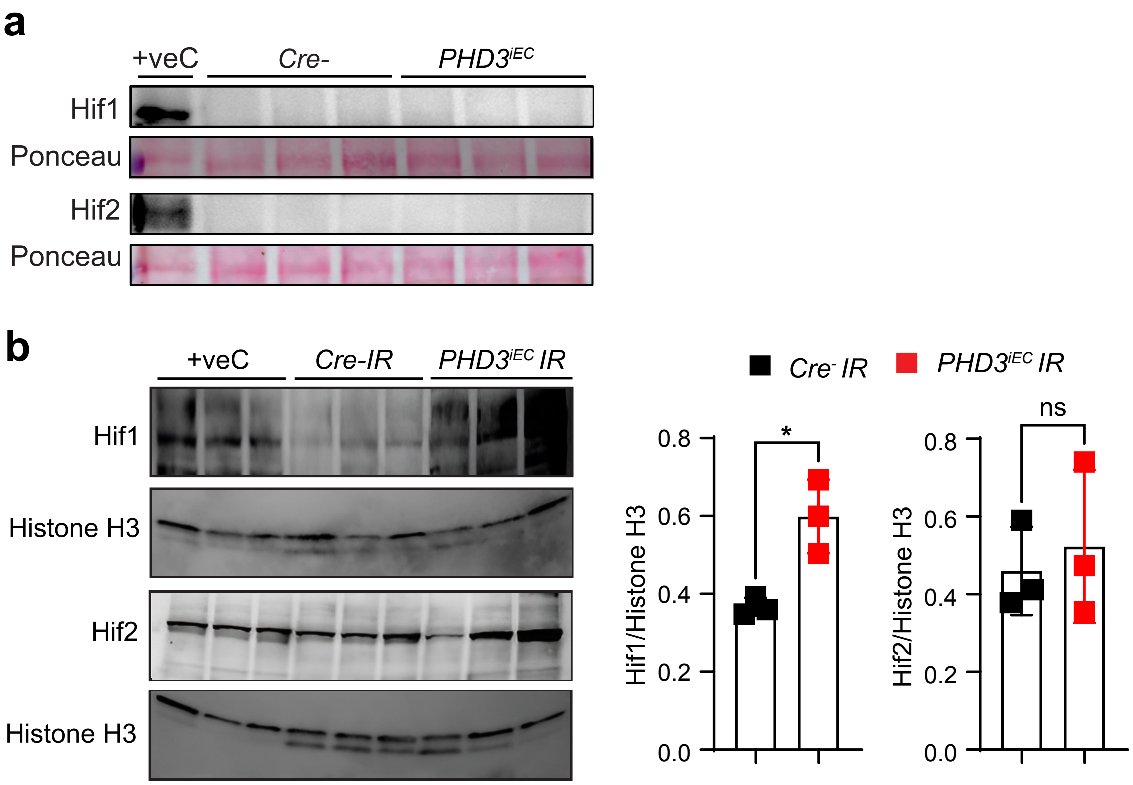

### Supplementry Figure 7

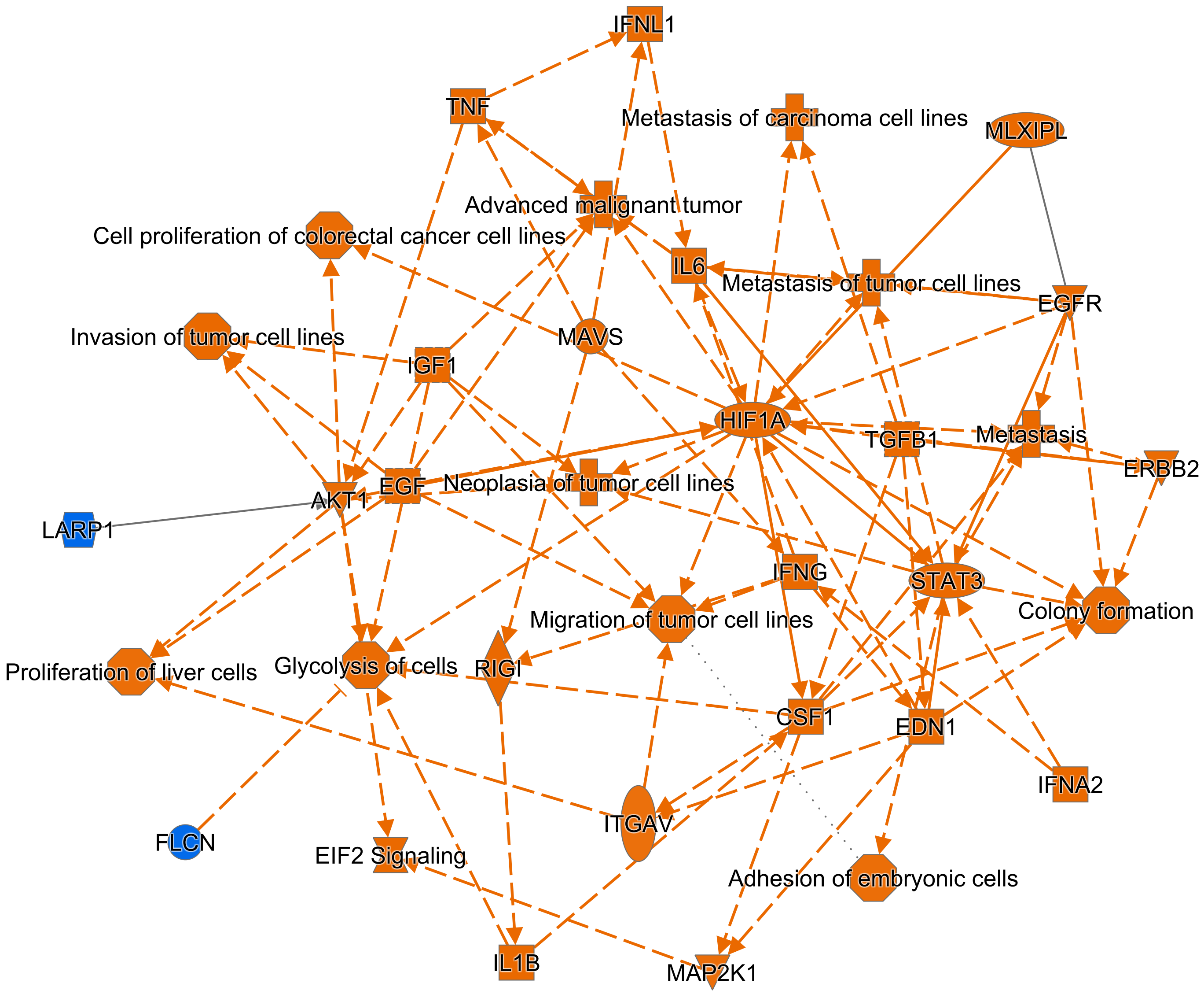

### SUPPLEMENTARY METHODS

#### Animals

Mice were maintained in a specific pathogen-free facility on a 12-hour light/12-hour dark cycle. Littermates were randomly assigned to the experimental groups. For the renal artery clamping procedure, anesthesia was administered via i.p. injection of xylazine (10 mg/kg) and ketamine (90–120 mg/kg). A small midline abdominal incision was made to access the left renal pedicle, which was clamped using a microaneurysm clamp for 25 min, while the right kidney remained untouched. Following the occlusion period, the clamp was released, and successful reperfusion was visually confirmed. The incision was closed using a 6-0 suture, and the skin was secured with Michel miniature clips. Body temperature was continuously monitored using a rectal probe and maintained at 37°C using a heating pad throughout the procedure. All experimental animals were included unless they showed abnormal behavior or infection post-procedure. Reporting followed the ARRIVE 2.0 guidelines [2].

#### RNA analysis by RT-PCR

RNA was extracted and used for real-time PCR analysis as previously described [3]. Mouse and human primer sequences are listed in Supplemental Table 1. RT-PCR was performed using the QuantStudio 3 Real-Time PCR system (Applied Biosystems) with SYBR Green or TaqMan PCR Master Mix. 18S rRNA was used for normalization.

#### Protein analysis by immunoblotting

Nuclear proteins were extracted using the NE-PER Nuclear and Cytoplasmic Extraction Reagent (Thermo Fisher Scientific). Nuclear protein extracts were separated by SDS-PAGE, transferred onto a nitrocellulose membrane, and incubated with either HIF-1α (Cayman; #10006421) or HIF-2α (Novus Biologicals; #NB100-122) antibodies at 4°C. After overnight incubation, the membrane was treated with a secondary antibody (Novus Biologicals), and chemiluminescent signals were detected using the SuperSignal West Femto Chemiluminescent Substrate (Thermo Fisher Scientific), followed by imaging with the iBright Imaging System (Thermo Fisher Scientific).

**Histopathological and Immunofluorescence analysis**

For histological analysis, kidneys were fixed in 10% formalin buffer and embedded in paraffin. To evaluate tubular damage and interstitial fibrosis, 5-μm transverse kidney sections were stained separately with hematoxylin and eosin (H&E) and Picro-Sirius red. Tubular injury was semi-quantitatively assessed by calculating the percentage of tubules in the corticomedullary junction that exhibited necrosis, brush border loss, cast formation, and tubular dilation (0, unaffected; 1:1–25%; 2:26–50%; 3:51–75%; 4:76–100%) [4]. Fibrosis was quantified as the percentage of the Picro-Sirius red positive area using ImageJ software (http://rsbweb.nih.gov/ij). For tubular injury scoring and fibrosis quantification, 10 and 5 random visual fields of the corticomedullary region per kidney section were analyzed at ×200 magnification, respectively.

For immunofluorescence staining, we used primary antibodies against PHD1 (Abcam; # ab113077), PHD2 (Novus Biologicals; # NB100-137), PHD3 (Novus Biologicals; # NB100-139), and Endomucin (Abcam; # ab106100). Goat anti-rat Alexa Fluor® 488 (Invitrogen; # A11006) and Goat anti-rabbit Alexa Fluor® 647 (Invitrogen, # A32733) were used as secondary antibodies. Sections were mounted with VECTASHIELD Vibrance Antifade Mounting Medium containing DAPI (Vector Laboratories, Inc. Catalog # NC1601055). Fluorescence images were acquired using a Nikon Ti2 widefield microscope and analyzed using the Fiji software (ImageJ). The Endomucin-positive area was quantified using ImageJ software.

**Flow Cytometry**

After perfusion with PBS, the kidneys were collected, finely chopped, and incubated at 37°C for 30 min in dissociation solution (Multi Tissue Dissociation Kit 2; Miltenyi Biotec; Cat. #130-110-203). The resulting cell suspension was sequentially filtered through 70 µm and 40 µm strainers, then mixed with 10 mL of cold staining buffer (PBS supplemented with 2% fetal bovine serum). After centrifugation at 400 g for 10 min, the cell pellets were treated with cold RBC lysis buffer, centrifuged again, and resuspended in staining buffer. Cells were then incubated with CD16/32 (Fc receptor blocker) and stained using various fluorophore-conjugated antibodies. The following antibodies were used: CD16/32 (eBioscience™, Catalog # 14-0161-85), CD45 (clone 30-F1, BioLegend; Catalog # 103140), F4/80 (clone BM8; BioLegend; Catalog # 123110), CD11b (clone M1/70, BioLegend; Catalog # 101216), CD3 (clone 17A2, BioLegend; Catalog # 100210), Ly6C (clone HK1.4; BioLegend; Catalog # 128010), and Ly6G (BD Biosciences; Catalog # 560600).

***Preparation of single-cell suspension***

Single-cell suspensions were generated using the Multi Tissue Dissociation Kit 2 (Miltenyi Biotec, Cat# 130-110-203), following the manufacturer's protocol. In brief, kidneys were sectioned into 6–8 pieces and placed in gentleMACS C tubes with 5 mL dissociation buffer (containing Buffer X, Enzyme P, Buffer Y, Enzyme D, and Enzyme A). Tissue dissociation was performed using the gentleMACS Dissociator with the Multi_E_01 program, followed by a 30-minute incubation at 37 °C with continuous rotation in a MACSmix Tube Rotator. The dissociation process was completed using the Multi_E_02 program. The reaction was halted by adding 10 mL of neutralizing buffer

(1X PBS with 2% fetal bovine serum), and the cell suspension was sequentially filtered through 100 µm and 30 µm strainers. After centrifugation at 400 g for 10 min, red blood cells were lysed with 1 mL of RBC lysis buffer on ice for 1 min, followed by neutralization and further centrifugation. Cells were washed, resuspended in 1X PBS with 0.04% BSA, and filtered through a 30 µm strainer. Cell counts and viability were assessed using a Nexcelom Cellometer Auto 2000 with AOPI fluorescent staining, yielding single-cell suspensions with over 80% viability.

**scRNA-seq data analysis**

The matrix files, which summarize the alignment results, were imported into Seurat (v5.1.0, Satija Lab, NYGC) and each individual sample was transformed to Seurat object. Only genes expressed in three or more cells and cells expressing at least 200 genes or more were used for further analysis. Cells with <500 unique molecular identifier (UMI) counts (cell fragments) and >100,000 UMI counts (potential cell duplets) were excluded. Furthermore, cells with a mitochondrial gene percentage of over 50% and low-complexity cells such as red blood cells with <0.8 log10 genes per UMI counts were also excluded from the analysis [5]. The merged data were normalized and scaled, and principal component analysis (PCA) was performed. The top 20 principal components were chosen as neighbors, and cell clustering was performed with a resolution of 0.1. Cell clusters were visualized in two-dimensional space using Uniform Manifold Approximation and Projection (UMAP). Both genotypes showed similar clustering and cell populations in separate and overlapping DimPlots. To identify cell clusters, marker genes were assessed using "FindAllmarkers" function of Seurat with the setting of min.pct of 0.25 and logfc.threshold of 0.25. Cell clusters were annotated based on the expression of top marker genes, as supported by published studies. Differential gene expression analysis was performed, and genes with adjusted $P<0.05$, $\log_2 FC > 0.2$ were considered significantly regulated. Significant genes (adjusted $P<0.05$) were used for gene set enrichment analysis using EnrichR (https://maayanlab.cloud/Enrichr/) and a bubble plot was created using R Studio.

#### ***Human snRNA-Seq data analysis***

Human snRNA-Seq datasets from AKI and control kidney tissues (17 AKI and 6 controls) were extracted from the Gene Expression Omnibus (GSE210622). Raw data were analyzed using Seurat's best-practice workflow for data integration using the reciprocal PCA approach (https://satijalab.org/seurat/articles/integration_rpca.html) with default parameters. Clusters were then analyzed for marker genes and EC cluster (*CD34*$^+$ and *EMCN*$^+$) was extracted, combined, and analyzed following Seurat's best practice standard workflow for data integration and analysis (https://satijalab.org/seurat/articles/integration_introduction.html). Differential gene expression analysis was performed, and significantly upregulated genes (adjusted $P<0.01$, $\text{Log}_2\text{FC}>0.25$) were subjected to gene network analysis using the IPA software (Qiagen).

#### **Cytokine Array**

To detect 111 mouse cytokines, kidney tissue homogenates were analyzed using a Proteome Profiler Mouse XL Cytokine Array Kit (R&D Systems # ARY028) according to the manufacturer's instructions. The signals were detected using a chemiluminescent iBright Imaging System (Thermo Fisher Scientific). Analyses were performed using ImageJ software.

#### **Cell Culture**

Human primary pulmonary artery EC (HPAEC) were obtained from ATCC and grown on gelatin-coated dishes in Endothelial Cell Basal Medium-2 (Lonza, Catalog # CC-3156) supplemented with EGM-2 SingleQuots Supplements (Lonza, Catalog # CC-4176). PPAEC were transfected using HiPerFect transfection reagent (Qiagen, Catalog # 301707) with 10 nM PHD3, ARNT, or Allstar negative control small interfering RNAs (siRNAs), all of which were purchased from Qiagen (FlexiTube GeneSolution PHD3; Catalog# SI03023083, ARNT Catalog# SI00304220). RT-PCR was used to assess transfection efficiency. After transfection, cells were analyzed under baseline conditions or following stimulation with 50 ng/ml IFN-γ (Sigma, Catalog# 117001) for 1.5hr.

**Lentiviral infection**

Control, EGLN3, and packaging plasmids were purchased from Addgene. pLp6.3-EglN3 and the control plasmid pLp6.3-Reverse were gifts from Susanne Schlisio (Addgene; #79115, #79117, #12260, and #12259). To generate lentivirus particles, HEK293T cells were co-transfected using jetPRIME transfection reagent (polyplus) with lentiviral vector (1µg/µl) and packaging plasmids as follows: psPAX2 (1µg/µl), pMD2.G (700ng/µl). Twenty-four- and 48-hours post-transfection, the supernatants containing lentiviral particles were collected and filtered through 0.45-µm filters. Lentiviral transduction of HPAECs was performed using lentiviral particles containing medium and polybrene (10µg/ml). After blasticidin selection, lentivirus-transduced cells were harvested and used for further experiments. Overexpression of PHD3 (EGLN3) was verified by immunocytochemistry.

**Supplemental Table 1.** Primer sequences. Shown are sequences of primers used for the expression analysis of the indicated mouse and human genes by RT-PCR.

| GENE | FORWARD PRIMER | REVERSE PRIMER |
|---|---|---|
| **Mouse** | | |
| *Loxl2* | 5′-GATCTTCAGCCCCGATGGA-3′ | 5′-CAAGGGTTGCTCTGGCTTGT-3′ |
| *Tgfb1* | 5′-TGGCGAGCCTTAGTTTGGA-3′ | 5′TCGACATGGAGCTGGTGAAA-3′ |
| *Acta2* | 5′CCTGACGCTGAAGTATCCGATA-3′ | 5′-TTTTCCATGTCGTCCCAGTTG-3′ |
| *Havcr1* | 5′-AAACCAGAGATTCCCACACG-3′ | 5′-GTCGTGGGTCTTCCTGTAGC-3′ |
| *F4/80* | 5′-CTTTGGCTATGGGCTTCCAGT-3′ | 5′GCAAGGAGGACAGAGTTTATCGG-3′ |
| *Cxcl10* | 5′-GATGACGGGCCAGTGAGAA-3′ | 5′-GCTCGCAGGGATGATTTCAA-3′ |
| **Human** | | |
| *PHD1* | 5′-GAATCAGAACTGGGACGTTAAGGT-3′ | 5′-CGGCCCTCAGGGAAGATC-3′ |
| *PHD2* | 5′-GCTTTGTTTGCCCCAGAGTATT-3′ | 5′-GAATGTCCCTCCCAATCCTTAAT-3′ |
| *PHD3* | 5′-GAGGCAATGGTGGCTTGCTA-3′ | 5′-CCACGTGGCGAACATAACC-3′ |
| *ICAM1* | 5´-CCACAGTCACCTATGGCAAC-3′ | 5′-AGTGTCTCCTGGCTCTGGTT-3′ |
| *VCAM1* | 5´-GCTTCAGGAGCTGAATACCC-3′ | 5´-AAGGATCACGACCATCTTCC-3′ |
| *CXCL9* | 5′-TCCACCTACAATCCTTGAAA-3′ | 5′-TGCTGAATCTGGGTTTAGAC-3′ |
| *CXCL10* | 5′-TGAATCAAACTGCCATTCTG-3′ | 5′-GTACAGCGTACAGTTCTAGA-3′ |
| *TAP1* | 5′-CAAGAGCCACAGGTATTTGG-3′ | 5′-ACTGCAGCAGCTGTGATTTC-3′ |
| *IFIT3* | 5′-GCTGAAGGAGAGCAGTTTGTTGA-3′ | 5′-AGGACATCTGTTTGGCAAGGA-3′ |
| *IFITM3* | 5′-GGCCCCAACCTGGGATT-3′ | 5′-AAATGTACCTTAGAGCAACATGCAA-3′ |
